# An extensive and unbiased genome-wide scan for parent-of-origin expressed genes in the pig clarifies the conservation landscape of genomic imprinting

**DOI:** 10.1101/2025.03.09.642261

**Authors:** Mathilde Perret, Nathalie Iannuccelli, Sophie Leroux, Katia Fève, Patrice Dehais, Eva Jacomet, Jean-Noël Hubert, Carole Iampietro, Céline Vandecasteele, Sarah Maman-Haddad, Thomas Faraut, Laurence Liaubet, Agnès Bonnet, Cécile Donnadieu, Juliette Riquet, Julie Demars

## Abstract

Genomic imprinting, a mechanism resulting in parent-of-origin expression of genes through epigenetic regulation, intersects with a broad range of biological fields including evolution, molecular genetics and epigenetics and determinism of complex traits. Although next generation sequencing technologies enable nowadays to detect imprinted genes in a genome-wide manner, a wide spectrum of this phenomena is evaluated only in humans and mice.

Here, we propose to map genes showing a parental expression bias in hypothalamus, muscle and placenta in piglets around birth using an extensive and unbiased strategy that relied on reciprocal crosses, genetics reconstruction of parental phases after imputation and statistical analyses discriminating parent-of-origin from allele-specific expression. We detected 440 unique genes with a weak to exclusive parental expression bias including 114 unique genes with an imbalance ratio above 25:75. About thirty imprinted genes are common to human and/or mice and an equivalent number is shared between tissues, suggesting an overall weak conservation landscape of genomic imprinting. Interestingly, we identified novel parent-of-origin expressed genes involved in neurodevelopmental (*PITRM1*, Pitrilysin Metallopeptidase 1) and fetal growth (*FAM20B*, Glycosaminoglycan Xylosylkinase and *POU6F2*, POU Class 6 Homeobox 2) functions. In addition, deeper analyses of specific loci likely highlighted lineage-specific imprinted genes such as a Zinc Finger Protein 300-like gene as well as specific imprinted isoforms of *COPG2* (COPI Coat Complex Subunit Gamma 2), a gene showing conflicting data in the literature.

Altogether, our results bring pig as the most comprehensively and exhaustively documented species for genomic imprinting after human and mice organisms. A weak conservation of this mechanism across species and tissues suggested a distinction between a small number of core imprinted genes and others parent-of-origin expressed genes that seemed subjected to evolutionary forces for acquiring their imprinting status either in a lineage-specific or tissue-specific manner.

## Background

Genomic imprinting is an original molecular phenomenon leading to a preferential allele-specific expression dependent on the parental origin. Genomic imprinting mechanism is a form of epigenetic regulation found in placental mammals and flowering plants (Wolf et al., 2014). Parent-of-Origin Expressed (POE) genes, often called imprinted genes, are found isolated or as clusters across the genome, representing 1% to 2% of the total gene content in the best studied mammals such as mice, rat and human species (Tucci et al., 2019). Imprinted genes include different types with approximately 1/3 being protein-coding genes and the rest being non-coding RNAs genes comprising long (lncRNAs) and small RNAs (microRNAs and snoRNAs) (Perez et al., 2016). The greatest variability of imprinting patterns (i.e. parental expressed allele exhibiting a weak to robust parent-of-origin effect between 45:55 and 10:90) was observed for protein-coding genes (Ho-Shing and Dulac, 2019).

Imprinted genes play critical roles in foetal and post-natal development and adult tissue function (Tucci et al., 2019) as supported by various human imprinting disorders (Monk et al., 2019) as well as mutations within imprinted genes associated with major agronomic phenotypes (Hubert et al., 2024b). Initial studies on genomic imprinting, mainly focused on key imprinted genes expressing exclusively one parental allele, suggested a certain level of conservation for the number of imprinted genes and their associated imprinting patterns between tissues and species (Bischoff et al., 2009; Magee et al., 2010; Park et al., 2011). Nowadays, multiple studies demonstrated differences between both tissues and development stages with the placenta and brain showing the highest frequency of imprinted genes (Babak et al., 2015; Daskeviciute et al., 2025; Ho-Shing and Dulac, 2019). These results are supported by Bonthuis et al. who highlighted that number of imprinted genes in the arcuate nucleus of the hypothalamus was 100% to 300% higher than somatic skeletal tissues (Bonthuis et al., 2022).

More and more studies are focusing on the evolution of genomic imprinting mechanisms across species. Comparison of genomic imprinting phenomenon through the phylogeny of eutherians brought novel insights on molecular mechanisms of imprinting acquisition by distinguishing canonical from non-canonical imprinting, the latter being exclusively identified in muridae so far (Richard Albert et al., 2023). While canonical imprinting is driven by DNA methylation, non-canonical imprinting, a form of oocyte-specific acquisition of genomic imprinting, is established by the apposition of H3K27me3 on retrovirus sequences including LTR (Long Terminal Repeats) (Inoue et al., 2017). This specific pattern results in maternally DNA methylated imprints that govern paternal-specific gene expression in extra- embryonic cells (Inoue et al., 2017). The most studied species still remain Human and models organisms such as Mouse and Rat and some specific results were dedicated to marsupials (Edwards et al., 2019; Stringer et al., 2014). However, genomic imprinting studies on other species including livestock were often restricted to a few orthologous genes (Ishihara et al., 2024; Magee et al., 2010; Park et al., 2011) leading to a gap of knowledge about genomic imprinting across evolution.

Next-generation sequencing technologies focused on RNA enabled the identification of imprinted genes in a genome-wide manner allowing the detection of a large spectrum of parental biases in transcripts from various biological samples. However, such novel approaches applied to genomic imprinting mechanisms required dedicated experimental designs and specific bioinformatics and statistical methods to avoid false positive results (Edwards et al., 2023). Despite rigourous and stringent analyses, the studies mostly failed to detect novel imprinted genes due to uncomplete genome annotations in particular for non-coding RNAs. However, recent data obtained in pigs highlighted lineage-specific imprinted genes such as *KBTBD6* (Kelch Repeat And BTB Domain Containing 6) (Ahn et al., 2023) and *ZNF791*-like (Zinc Finger Protein 791) (Ahn et al., 2025) reinforcing the necessity to acquire depeer knowledge on genomic imprinting mechanisms in different mammals to better understand the emergence of imprinting during evolution on one hand and to improve our understanding of the role of imprinted genes for the construction of traits of interest in domestic species on the other hand.

Here, we propose to map imprinted genes in hypothalamus, muscle and placenta in piglets around birth using an exhaustive and unbiased strategy relying on reciprocal crosses, the genetic reconstruction of parental phases after imputation and an objective discrimination of POE from allele-specific expression. Genome-wide comparison of both number and distribution of imprinted genes along the pig genome showed a weak conservation of them across tissues and species (pig *vs.* human and mouse species) but, when conserved, exhibit a strong conservation of their imprinting patterns (i.e. the direction of parental expressed allele). These results suggest the distinction across evolution between a small number of core imprinted genes with shared POE and others that acquire their imprinting status in a lineage-specific manner. In addition, deeper analyses of specific loci likely highlighted novel imprinted genes including a *ZNF300-like* gene (Zinc Finger Protein 300) (*LOC100520903*) in pigs as well as specific imprinted isoforms of *COPG2* (COPI Coat Complex Subunit Gamma 2), a gene showing conflicting data in the literature.

## Results

### An exhaustive and unbiased strategy to detect parental expression bias

In order to detect parental expression bias, we relied on 2 different datasets generated from reciprocal crosses between two genetically distant pig breeds, the European Large White (LW) breed and the Asian Meishan (MS) breed (Fig. 1A). Piglets from the first dataset have been produced to highlight parental expression bias in muscle and hypothalamus at one day and the second dataset was dedicated to target imprinted genes in the placenta at 110 days of gestation.

**Figure 1:**
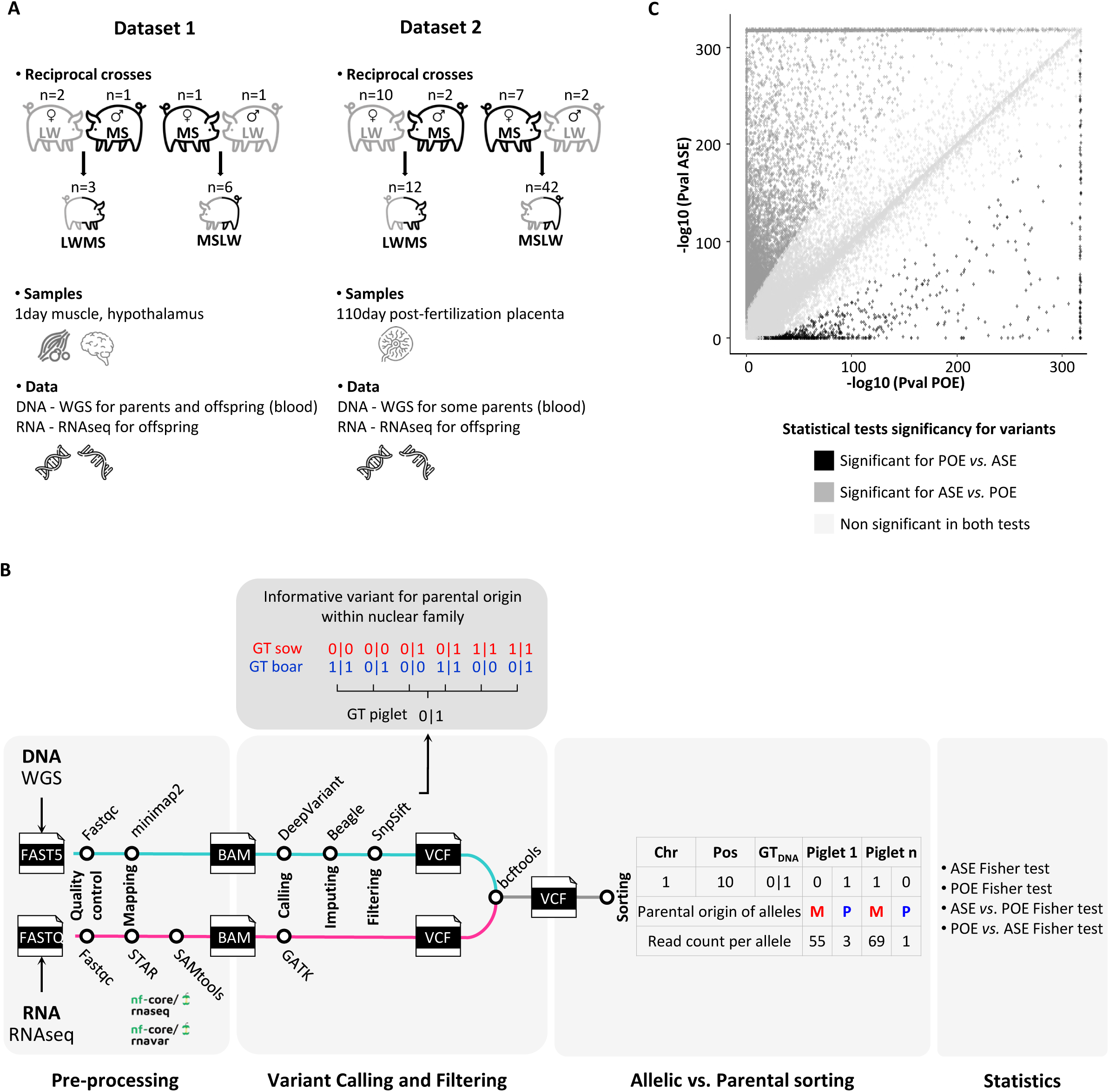
Experimental designs and approach to detect parent-of-origin expression. **A.** Large White (LW) pigs and Meishan (MS) pigs underwent reciprocal crosses at two different developmental stages (1 day after birth, dataset 1 and 110days post-fertilization, dataset 2). Tissue sample size and sequencing contents across both datasets are mentioned. **B.** Roadmap of preprocessing and statistical analyses leading to an unbiaised an extensive strategy to discriminate allele-specific expression (ASE) from parent-of-origin expression (POE). **C.** Scatter plot of statistical analyses for all informative variants in the three tissues (hypothalamus, muscle and placenta). For each variant, the values of -log10 (Pval ASE) and -log10 (Pval POE) from one sided Fisher exact tests have been plotted and the direction of the effect (ASE or POE) has been colored in black for the POE *vs*. ASE significancy, in dark grey for the ASE *vs.* POE Fisher test significancy and in light grey when it was not possible to discrminate between both directions. Only variants in black showing parental expression bias have been conserved for further analyses.

We developed an agnostic strategy to highlight parental-expressed genes since we explored the whole genetic variability of variants not taking into account the annotation of the pig reference genome at first glance (Fig. 1B). Calling of variants from complete genomic data followed by imputation allowed the detection of 28,863,478 and 32,431,979 variants in the first and second dataset, respectively. Filtering of variants has been performed to keep all informative heterozygous variants in offspring, those for which parental origin was traceable, and not exclusively those fixed in each breed as usually done. Thus, six possible parental genomic combinations enabled us to identify 24,024,152 parental informative variants in the hypothalamus and muscle and 13,025,625 in the placenta (Fig. 1B). In the meantime, calling of variants from transcriptomic data enabled us to identify 3,572,855, 2,204,937 and 792,903 variants in the hypothalamus, muscle and placenta respectively (Fig. 1B). Intersection of both lists of variants from genomic and transcriptomic data per tissue was performed to obtain position for which the parental expression bias could be exploited. From the intersection, we only filtered bi-allelic genomic positions comprising at least one individual per crossing direction, and for which the total number of reads was >= 15 in the hypothalamus and muscle, and >= 90 in the placenta, due to the 6 times greater number of F1s compared with the other two datasets. A total of 539,555, 322,599 and 191,901 variants were considered for further analyses in hypothalamus, muscle and placenta, respectively (Fig. 1B).

Different statistical analyses have been performed to discriminate Parent-of-Origin Expression (POE) from Allele-Specific Expression (ASE). We firstly assessed POE and ASE expression using Fisher tests at each genomic position in the three tissues. We also assessed which of the two mono-allelic expression (POE: POE *vs.* ASE or ASE: ASE *vs.* POE) was most probable, also using Fisher’s tests. From the 539,555, 322,599 and 191,901 variants analyzed in the hypothalamus, muscle and placenta, respectively, bi-allelic expression was concluded for a large proportion since neither ASE nor POE could be significantly determined for 531,357, 317,735 and 186,847 in the hypothalamus, muscle and placenta, respectively. We determined a set of 7748, 4597, and 4595 variants (Bonferroni correction, alpha = 0.01) in the hypothalamus, muscle and placenta for which the ASE was significantly more probable than POE (ASE *vs.* POE, Fisher test). Finally, we identified 450, 267, and 459 variants (Bonferroni correction, alpha = 0.01) for which the POE phenomenon was significantly more likely than ASE (POE *vs.* ASE, Fisher test) (Fig. 1C).

### Identification of parent-of-origin expressed genes, from weak imbalance to imprinting

Our strategy to identify parental expression bias in a genome wide manner relied on an agnostic approach : all variants were firstly considered since no filter were applied neither on the selection of variants from their original breed nor on the annotation of variants to maximize the possibility of identifying novel candidate genes for imprinting. We analyzed both the genomic distribution and the parental origin of variants previously detected to show POE (Fig. 2A and Additional file 1). While some variants from common parental-origin are grouped in clusters, others are more widespread suggested that one gene could be either targeted by several variants or a single variant. Annotation of significant variants was performed and we identified 174, 78 and 239 genes in the hypothalamus, muscle and placenta, respectively, including three unannotated variants, from the 450, 267, and 459 variants showing a significant POE. In total, this represented 440 unique genes with a significant parental expression bias. Several highly significant variants from the same parental origin were identified in well-known imprinted genes such as *IGF2 (Insulin Growth Factor 2)*, *MEG3* (*Maternally Expressed Gene 3*) and *MEST* (*Mesoderm Specific Transcript*) validating the appropriateness of our approach (Fig. 2A and Additional file 1).

**Figure 2:**
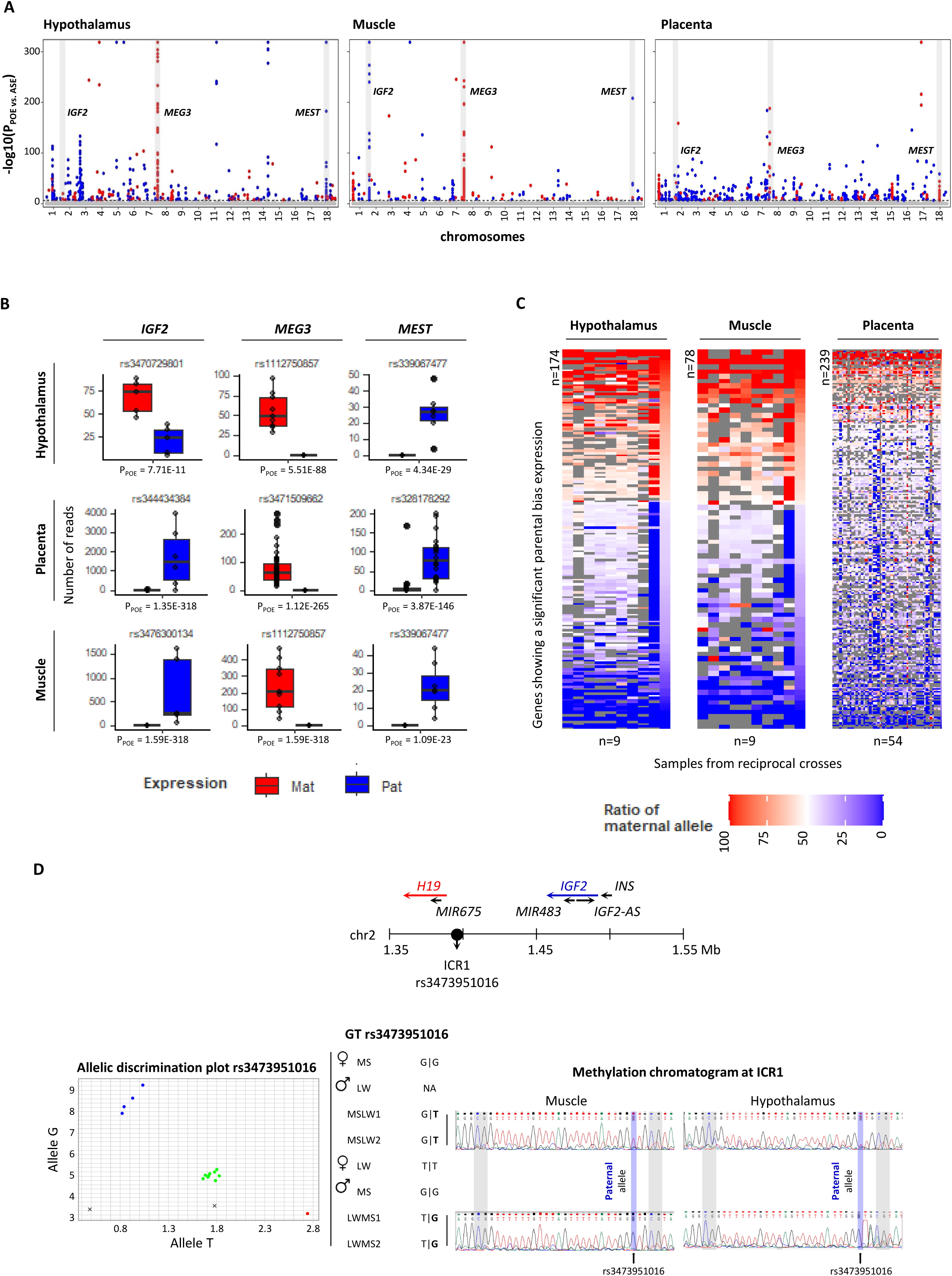
Genome-wide detection of parental expression bias in piglets around birth. **A.** Distribution of significant variants for POE *vs.* ASE statistical Fisher test along pig autosomes for each tissue. Each variant has been colored based on its major parental expressed allele in blue or red for paternal and maternal alleles, respectively. Some well-known imprinted genes such as *IGF2*, *MEG3* and *MEST* in pigs have been highlighted (grey boxes). **B.** Distribution of parental reads for three imprinted genes in each tissue. For each tissue, one significant variant tagging *IGF2*, *MEG3* and *MEST* genes has been considered and number of parental reads for informative individuals has been plotted. Reads from paternal and maternal origins have been colored in blue and red, respectively. **C.** Genes showing parent-of-origin expression. Each signficant variants for POE *vs.* ASE statistical Fisher test has been annotated using both Ensembl annotation release 113 and NCBI annotation release 106. When several variants tagged the same gene, the mean of POE has been considered leading to 174, 78 and 239 unique genes with parental expression bias in hypothalamus, muscle and placenta, respectively. The ratio of maternal allele has been plotted on the heatmap with blue corresponding to paternal bias expression and red corresponding to maternal bias expression. **D.** Parental methylation evaluation at the Imprinting Control Center 1 (ICR1) of the *IGF2* locus. The variant rs3473951016, located within ICR1, was informative in two nuclear families from reciprocal crossed of dataset 1 with the T allele coming from the boar in the first MSLW cross and the G allele coming from the boar in the second LWMS cross. Chromatograms issued form methylation sensitive PCR followed by Sanger sequencing of PCR products in hypothalamus and muscle showed that only paternal allele is methylated in both tissues.

For each gene identified, we analyzed more accurately the parental expression bias in the three tissues to catch different patterns (Fig. 2B). While much more genes showed a trend for the paternal expressed allele in the placenta, it was more balanced in both hypothalamus and muscle tissues. In the hypothalamus, muscle and placenta we observed 104, 47 and 195 genes showing a preferential paternal expression and 70, 31 and 44 genes with maternal expression bias respectively. Analysis of these ratios enabled us to split the identified genes into two expression subgroups : genes showing a weak to moderate parental expression bias (ratios ranging from 46:54 to 25:75 with a wide POE significancy values from Pval = 1.48E-10 to Pval = 1.35E-318), and genes showing strong parental bias to exclusive parental expression (ratios above 25:75 to 0:100 and Pval POE included Pval = 2.34E-09 for very low expressed genes and Pval = 2.66E-318, the limit of detection) (Fig. 2B and Additional file 1). Only 36%, 47% and 16% showed a strong magnitude of parental imbalance among the 174, 78 and 239 genes with POE in the hypothalamus, muscle and placenta, respectively. This represented 114 unique genes including 11 genes that did not correspond to a HUGO annotation but were annotated to genes of uncertain function (LOC symbols) from NCBI annotation release 106 (Additional file 1).

Analyses of parental reads distribution for *IGF2*, *MEG3* and *MEST* clearly highlighted an exclusive expression from the paternal allele for *MEST* in contrary to *MEG3* that was exclusively expressed from the maternal allele (Fig. 2C). The conclusion seemed as sharp for *IGF2* in the muscle and placenta with a full paternal expression but a switch to a maternal expression in the hypothalamus has been observed. These results are independent of the gene expression. We wished to address whether these differences in IGF2 expression patterns between tissues could be caused by a difference in parental origin of the methylation within the Imprinting Control Region 1 (ICR1), responsible for its imprinted state regulation. We molecularly characterised the parental origin of the ICR1 methylation (Fig 2D) using the rs3473951016 marker located within this region and informative in reciprocal crosses of the first dataset. In each cross direction, we observed the presence of the paternal allele in both tissues as shown on chromatograms of Fig. 2D. Thus, the *IGF2* gene, which showed a maternal bias of expression in the hypothalamus and an exclusive paternal expression in the muscle, seemed regulated by paternal methylation in both tissues at ICR1 (Fig 2.D).

### A weak conservation of imprinting genes across mammals in contrast to their imprinting patterns

In order to evaluate the conservation landscape of parental expression bias, we compared genes identified in piglets regardless the tissue analyzed to human and mouse species for which the characterization of imprinting is the most exhaustive. A total of 440 unique genes have been detected from the hypothalamus, muscle and placenta in piglets altogether (Fig. 3A) but only 114 of them showed a strong magnitude of parental imbalance above ratio of 25:75 (Additional files 1 and 2). The distribution along the pig autosomes of these genes as well as the position of mouse and human orthologous imprinted genes known from the literature is showed on Fig. 3A (with the 440 pig genes) and Additional file 2 (with only the 114 pig genes with strong parental imbalance). Only a small proportion of locations overlapped between species since only 25 genes with a parental expression bias were commonly identified in at least two species. All of these conserved genes across piglets, mouse and human were located in clusters and often showed an exclusive parental expression that is typical of known imprinted genes. For an additional dozen of common positions between species, it occurred that neighboured orthologous genes showed parental expression bias likely suggesting a smooth shift to acquire imprinting across evolution of genomes (Fig. 3A and Additional file 3).

**Figure 3:**
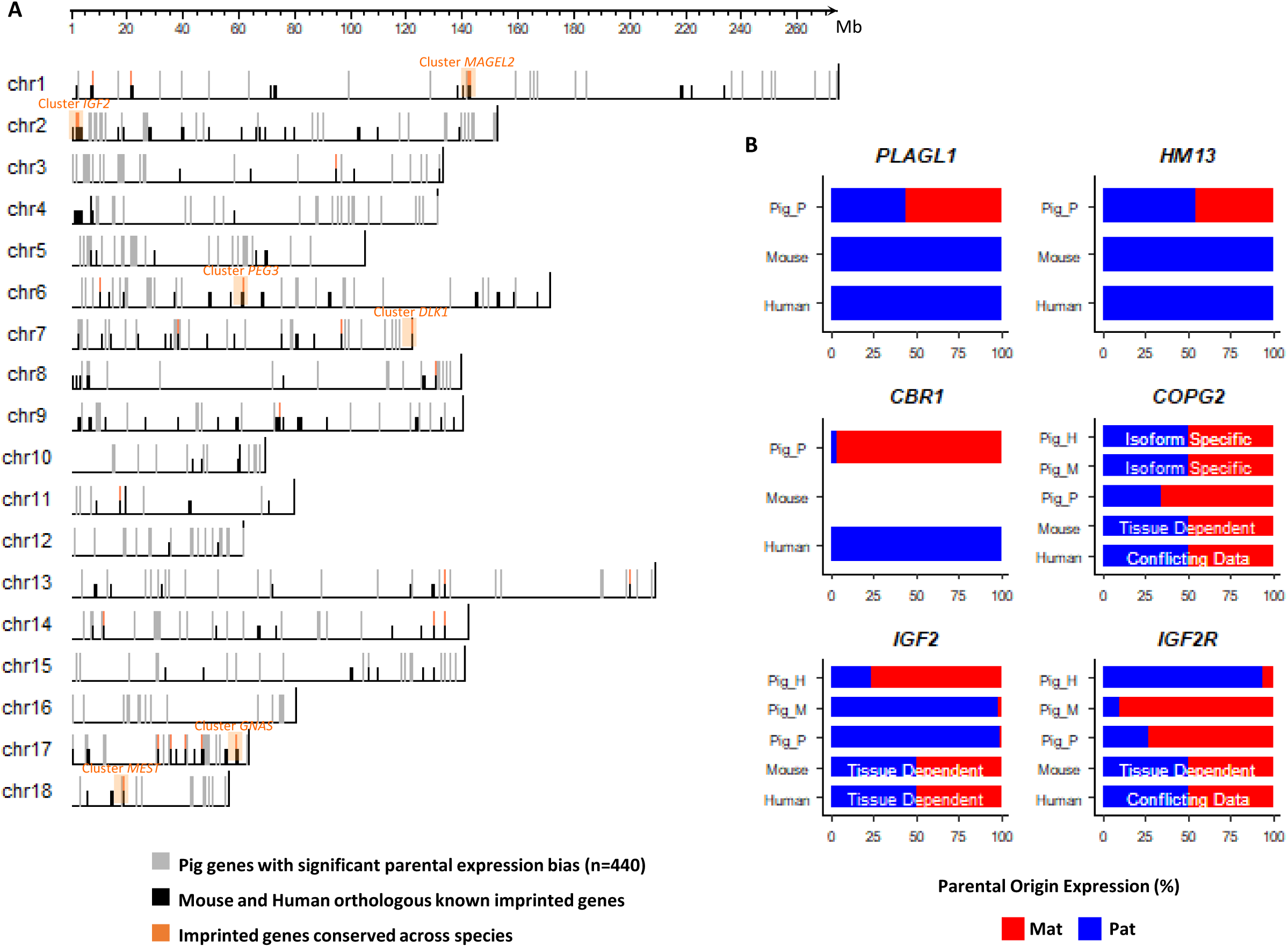
Conservation across human and mouse species of parent-of-origin expressed genes. **A.** Distribution along the pig autosomes of genes showing POE in piglets and orthologous imprinted genes identified iun both human and mouse species. The position of unique genes (n=440) from the 178, 74 and 239 identified in hypothalamus, muscle and placenta, respectively tissu has been plotted on the ideogram in light grey. Positions of mouse and human orthologous imprinted genes (Tucci et al., 2019) have been mentioned in black and overlapping of genes and positions between piglets, mouse and human, suggesting conservation across species, have been highlighted in orange. **B.** Parental expression bias in conserved parent-of-origin expressed genes. Among the few conserved imprinted genes, six showed contrasting imprinting patterns between piglets and mouse or human species. Parental expressed alleles are in blue and red for paternal and maternal, respectively. Pig_H : expression in hypothalamus of 1day piglets, Pig_M : expression in muscle of 1 day pigles, Pig_P : expression in placenta of 110 days piglet fetuses. Isoform specific corresponds to the parental expression of a specific-isoform.

In contrary, imprinting patterns (i.e., the direction of parental expressed allele) for the 25 imprinted genes are very well conserved except for 6 of them, which are visualized on Fig. 3B. Some discrepancies due to conflicting data or tissue dependency were already mentioned in the literature as suggested for *COPG2 (COPI Coat Complex Subunit Gamma 2)*, *IGF2* and *IGF2R (Insulin Growth Factor 2 Receptor)*. The three remaining genes with different imprinting patterns between species, including *PLAGL1 (PLAG1 Like Zinc Finger 1)*, *HM13 (Histocompatibility Minor 13)* and *CBR1 (Carbonyl Reductase 1)*, showed parental expression bias only in the placenta. These results might likely be explained *via* a specific placentation for pig *vs.* human and mouse presenting an epithelo-chorial and hemo-chorial placentation, respectively.

### The COPG2 locus, detection of the paternal COPG2IT1 gene and isoform-specific imprinting of COPG2

We tried to understand inconsistencies obtained for *COPG2* in piglets since discrepancies were also observed in human and mouse fro this gene. Indeed, several variants spanning the *COPG2* gene brought conflicting information from the parental expression bias with some variants suggesting bi-allelic expression, while other a trend for the maternal allele and others an exclusive paternal expression. A deep visual analysis of reads through the Integrative Genome Viewer (IGV) combined to the identification of isoforms from our transcriptomic data using Stringtie (Pertea et al., 2015), allowed to detect an novel antisens transcript in the hypothalamus that was not annotated so far in the pig genome (Fig. 4A). This antisens located between *COPG2* and *MEST* genes likely corresponding to the *COPG2IT1* gene that clearly showed a paternal expression based on parental reads distribution from rs3473698674, rs3473779585 and rs3475297787 (Fig. 4A).

**Figure 4:**
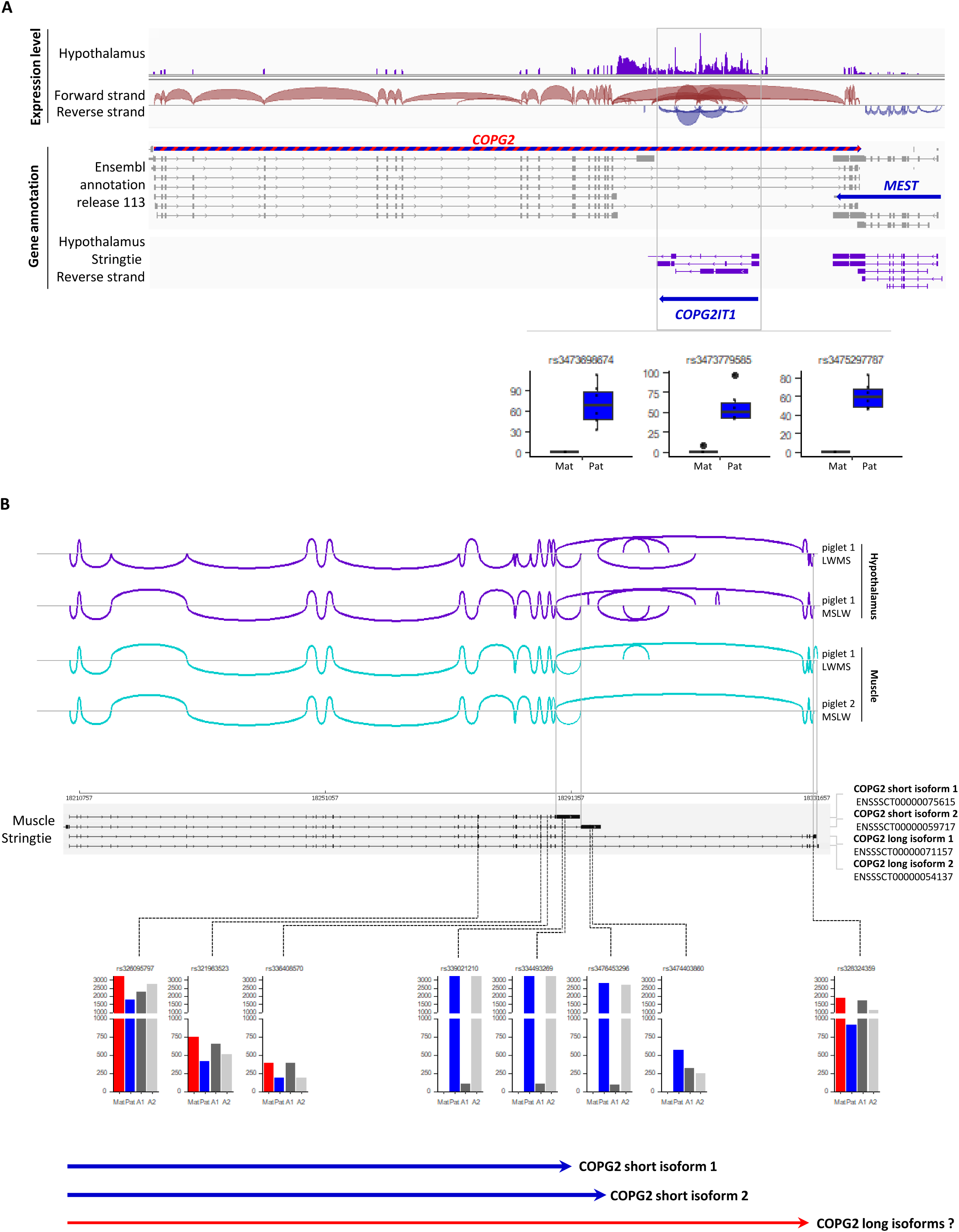
The *COPG2* locus, detection of paternal *COPG2IT1* and COPG2 paternal isoforms. **A.** Identification of the paternal *COPG2IT1* gene in hypothalamus. Screenshot of the *COPG2* locus using the Integrative Genome Viewer (IGV (Robinson et al., 2023)). Expression level in hypothalamus in one individual showed high coverage (grey box) from the reverse strand while none gene was located and annotated in the last available Ensembl annotation release 113. Transcript asssembly using Stringtie (Pertea et al., 2015) detected an antisens transcript at this location. Three significant variants fro POE *vs.* ASE statistical Fisher test (rs3473698674, rs3473779585 and rs3475297787) overlapped this novel transcript and showed an exclusive paternal expressed allele in hypothalamus strongly suggested the identification of *COPG2IT1* gene. **B.** Identification of paternal COPG2 isoforms. The COPG2 transcripts showed short and long isoforms in both hypothalamus and muscle tissues. Transcript assembly using Stringtie (Pertea et al., 2015) improved 3’ UTR of shortest isoforms compared to Ensembl annotation release 113. Analyses of POE of variants spanning *COPG2* gene showed that variants rs339021210, rs3344933269, rs3476453296 and rs3474403860 tagged exclusively last exons of COPG2 shortest isoforms. Only paternal allele is expressed for all of them, suggesting a paternal expression of shortest isoforms of COPG2. None informative and significant variants tagged exclusively longest isoforms of COPG2, making their expression patterns unconclusive.

Beside the identification of the COPG2IT1 non-coding transcript within the *COPG2* locus, visual and manual inspection of the *COPG2* gene itself highlighted the parental expression for specific COPG2 isoforms (Fig. 4B). Annotation of *COPG2* isoforms from Ensembl annotation release 113 and NCBI annotation release 106 suggested short and long isoforms sharing the first exons for all of them. The reconstruction of transcripts that were present in hypothalamus and muscle allowed to detect at least four distinct isoforms, the two shortest (ENSSSCT00000075615 and ENSSSCT00000059717) and two longest isoforms (ENSSSCT00000071157 and ENSSSCT00000054137) as showed on sashimi plots in Fig. 4B. Based on the location of informative variants within the different exons of *COPG2* gene, we observed a sharp switch between variants that were common to both shortest and longest isoforms (rs326095797, rs321963523, rs336408570 and rs319794796) and variants specific of shortest isoforms (rs339021210 and rs334493269 for COPG2 short isoform 1 and rs3476453296 and rs3474403860 for COPG2 short isoform 2) Fig. 3C. We concluded that both shortest isoforms seemed exclusively expressed from the paternal allele while it was more difficult to conlude for the longest isoforms between maternal or bi-allelic expression since some exons are shared with the downstream *MEST* exons.

### Identification of novel imprinted genes shared between tissues and likely lineage-specific

In order to better characterize imprinted genes in piglets and further explore genes showing parental expression bias based on statistical tests, we evaluated their repartition between the three tissues. Among the 440 unique genes identified across tissues, the majority seemed specific of one tissue with 136, 44 and 221 out of 174, 78 and 239 for hypothalamus, muscle and placenta, respectively (Fig. 5A). While 24 genes were shared between hypothalamus and muscle, only 7 were common between hypothalamus and placenta although these tissues carry the largest number of genes with POE suggesting the specificity of these tissues for genomic imprinting mechanisms. We highlighted 6 genes including *IGF2*, *IGF2R*, *MEG3*, *MEST*, *KBTBD6* and *FAM20B (FAM20B Glycosaminoglycan Xylosylkinase)* that showed a bias of parental expression in the three tissues (Fig. 5A and 5B).

**Figure 5:**
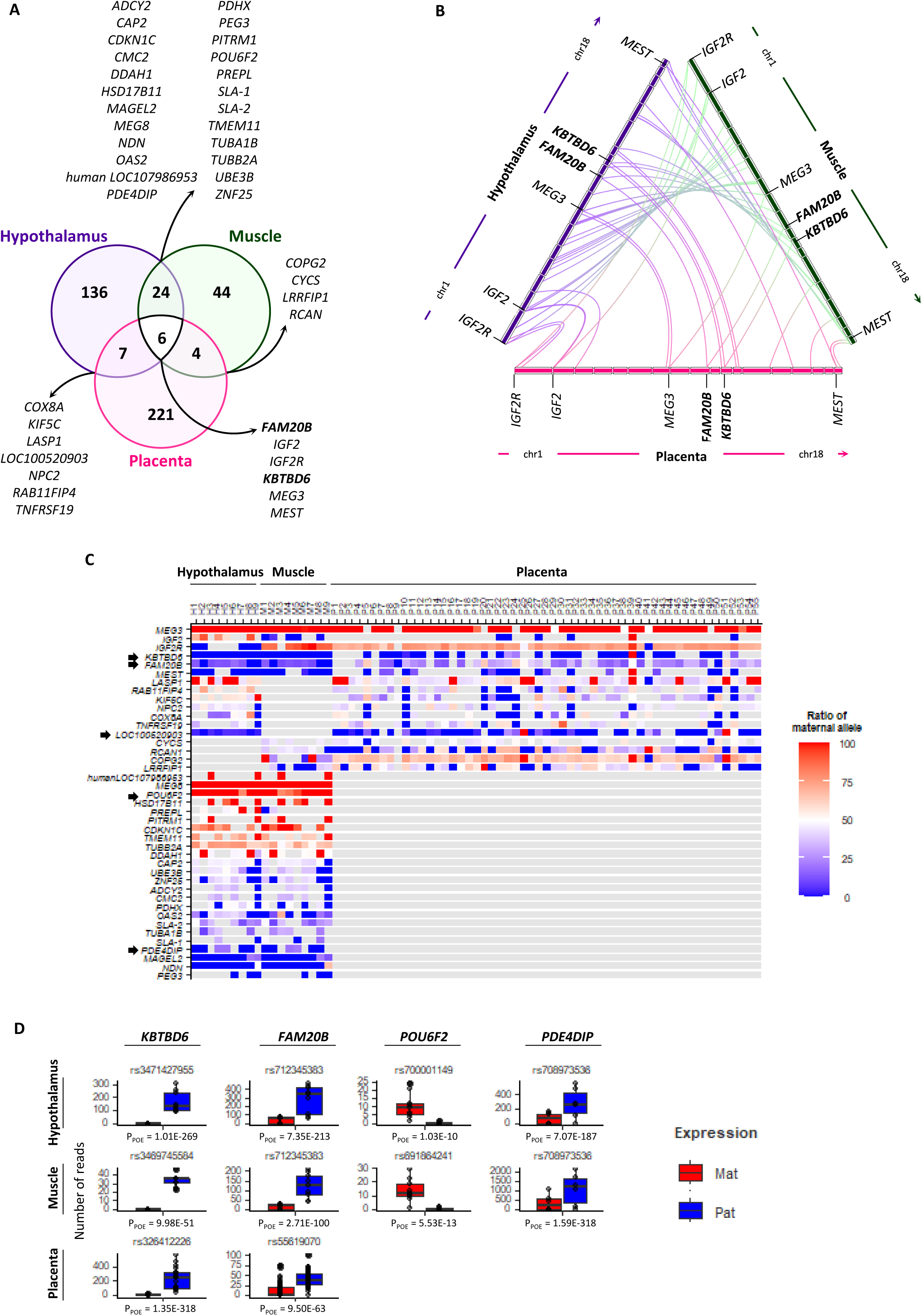
Conservation across tissues of parent-of-origin expressed genes in piglets. **A.** Venn diagram of parent-of origin expressed genes. A venn diagram was build from the 174, 78 and 239 genes with a significant parental expression bias in hypothalamus, muscle and placenta, respectively. Most of detected genes seemed tissue-specific. **B.** Distribution across pig chromosomes of parent-of-origin expressed genes detected in at least two tissues. A focus on the six shared genes is mentioned. **C.** Heatmap of POE focused on genes detected in at least two tissues. The ratio of maternal allele has been plotted on the heatmap with blue corresponding to paternal bias expression and red corresponding to maternal bias expression. **D.** Parental expression bias of novel imprinted genes. Among genes showing POE in at least two tissues, genes with a strong or exclusive parental imbalance (above 25 : 75) and unknwon so far for imprinting were selected. For each tissue, one significant variant tagging *KBTBD6*, *FAM20B*, *POU6F2* and *PDE4DIP* genes has been considered and number of parental reads for informative individuals has been plotted. Reads from paternal and maternal origins have been colored in blue and red, respectively.

Further analyses were then conducted on the genes shared between at least two tissues (n=41). The magnitude of parental expression bias was also analysed in greater detail to detect novel imprinted genes (Fig. 5C and Additional files 1 and 2). Most of genes showed a weak to moderate ratio of one parental allele except for *MEG3*, *IGF2R*, *KBTBD6*, *FAM20B*, *LOC100520903 (ZNF300-like)*, *MEG8 (Maternally Expressed 8, Small Nucleolar RNA Host Gene)*, *POU6F2 (POU Class 6 Homeobox 2)*, *PDE4DIP (Phosphodiesterase 4D Interacting Protein)*, *MAGEL2 (MAGE Family Member L2)* and *NDN (Necdin, MAGE Family Member)* for which the parental imbalance ratio is above 10:90 suggesting a typical pattern of imprinting (Additional file 1). Some of these genes are well-known imprinted genes while others have never been described before such as *FAM20B*, *LOC100520903*, *POU6F2* and *PDE4DIP* (Fig. 5D). An exclusive paternal expression has been observed for *KBTBD6* and an exclusive maternal expression has been shown for *POU6F2*. Although the pattern of imprinting for *FAM20B* and *PDE4DIP* was not restricted to a single parental allele, a high significant bias was obtained in direction of the paternal allele (Fig. 5D). In addition, a focused analysis on the unannotated *LOC100520903* gene, a zinc finger protein 300-like gene that seemed specific to the pig species, was carried out. The *ZNF300-like* gene located between *RNF216 (Ring Finger Protein 2016)* and *OR10AH1 (Olfactory Receptor Family 10 subfamily AH member 1)* showed epigenetic regulation marks from FAANG public dataportal (https://data.faang.org/home) including CpG island, CTCF binding site, imprinting marks such as H3K4me3 spanning upstream the expected transcription start site strongly suggesting its expression and putatively an imprinting pattern (Fig. 6A). This region seems to have undergone different evolutionary forces as observed on the dot plot between several species demonstrating the acquisition of complexity in orthologous regions across speciation through an expansion of highly repeated elements of Type I transposons family (Fig. 6B). In piglets, this novel gene showed an exclusive paternal expression in both hypothalamus and placenta (Fig. 6C).

**Figure 6:**
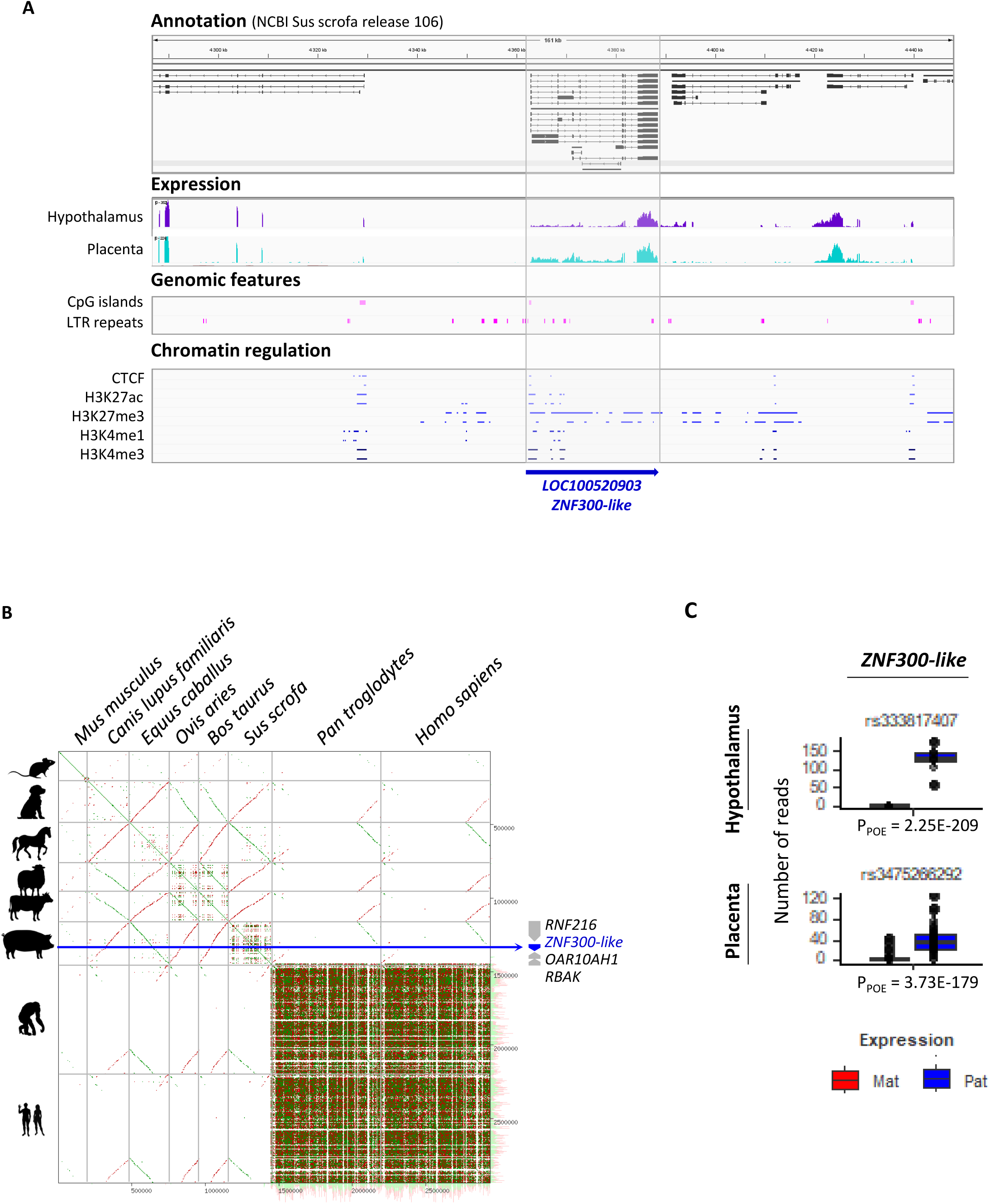
The paternal *ZNF300-like* gene (*LOC100520903)* ; a putative lineage-specific imprinted gene. **A.** Detection of the *LOC100520903*, a ZNF300-like gene, in both hypothalamus and placenta. Screenshot of the *RNF216-RBAK* locus using the Integrative Genome Viewer (IGV (Robinson et al., 2023)). Expression level in hypothalamus in one individual showed coverage from the forward strand of the annotated *LOC100520903* only in the NCBI Sus scrofa release 106. Distribution of CpG island, LTR repeats and chromatin marks regulation along the region using publicaly available FAANG dataset (https://data.faang.org/home). **B.** Comparison of sequences of the *RNF216-RBAK* region in several species. Fasta sequences of *RNF216-RBAK* region from mouse (GCA_000001635.9), dog (GCA_011100685.1), horse (GCA_041296265.1), cow (GCA_002263795.4), sheep (GCA_016772045.2), pig (GCA_000003025.6), chimpanzee (GCA_028858775.2) and human (GCA_000001405.29) were aligned and compared across each other. The dot plot showed the evolution of the region through expansion of repeated elements across speciation. **C.** Parental expression bias of the *ZNF300-like* gene. For hypothalamus and placenta tissues, one significant variant tagging the *LOC100520903* gene has been considered and number of parental reads for informative individuals has been plotted. Reads from paternal and maternal origins have been colored in blue and red, respectively.

## Discussion

We provided the results of a large-scale analysis of POE in piglets around birth aimed at minimizing any bias, whether at the experimental design (reciprocal crosses) or downstream analysis level (informativity and annotation of variants), as per latest recommendations (Edwards et al., 2023). Our strategy is based on reciprocal crosses between two breeds with very high genetic heterogeneity to maximize heterozygosity in the offspring and to discriminate between allele-specific and parent-specific expression. The use of parthenogenotes embryos, that are experimentally manipulated, as well as considering only gynogenotes embryos without comparison to androgenotes embryos, does not allow this distinction to be made and therefore seems less accurate for genome-wide mapping of genes subject to genomic imprinting (Ahn et al., 2025, 2023). In addition, the acquisition of genomic data in both parents and offspring allow to exploit the whole genetic variability by selecting all the heterozygous variants in offspring for which the parental origin can be deduced. Farm animal populations are polymorphic unlike congenic mouse lines. Although it is possible to select only the fixed variants in each breed to mimic the approach used for reciprocal crosses in mice, this is much more restrictive (Andergassen et al., 2017; Oczkowicz et al., 2018; Quan et al., 2024; Wu et al., 2020). Here, we were able to exploit around 24 million informative variants from genomic data, whereas most other studies analyzing only breed-specific variants from reciprocal crosses exploit around 5 times fewer variants (Quan et al., 2024). However, the lack of genetic informativity still remains the greatest source of incompleteness in the detection of imprinted genes, as was the case in our study where we were unable to identify the *PEG10* gene that is a well-known imprinted gene in pigs (Ahn et al., 2020).

Furthermore, all the available studies use variant annotation as an additional filter before statistical analysis in order to consider only annotated genes. This assume that the genomes are perfectly annotated, which is not the case, particularly for non-coding genes representing 2/3 of the imprinted genes identified to date (Perez et al., 2015). Their identification in mammalian genomes is difficult and are often poorly annotated (Mattick et al., 2023). For example, *KCNQ1OT1* and *NESP-AS*, known to regulate imprinting at the *CDN1C-KCNQ1* loci and *GNAS* in mammals respectively, do not appear in the porcine genome annotation (Ensembl annotation release 113 and NCBI annotation release 106 databases). Here, we chose to apply annotation filter *a posteriori* in order not to restrict the list of significant variants *a priori*. Several variants did not appear to be located in annotated genes or were poorly annotated, and manual inspection of these variants often revealed a gene, as was the case for the *COPG2IT1* gene. The detection of such regions demonstrates the power of an agnostic approach. Altogether, we detected 440 unique genes with POE within three tissues in piglets including 114 with a strong to exclusive parental bias above the ratio 25:75 while the most robust and extensive approach to date identified 179 genes with POE and only 17 had a 5:95 average expression bias compared to 39 in our study demonstrating the power of our strategy (Quan et al., 2024).

The use of two datasets of very different sizes demonstrated that the power of a larger setup allowed to detect much more precise parental expression biases (ratio 45:55) rather than a larger number of genes with very strong or even exclusive expression biases. Among the 114 genes showing a strong magnitude of POE imbalance, 65% and 35% showed paternal and maternal expression, respectively. These results were consistent with the data available in model species for imprinting such as humans, rats and mice (https://www.geneimprint.com/), contrary to what has been observed in the literature in pigs, where these ratios appear to be balanced between the 2 parental origins (Quan et al., 2024). Furthermore, the characterization of the type of genes showed that a large majority of these genes are protein-coding genes (more than 75%) in our study in contrary to the existing knowledge in mice (Perez et al., 2015). In addition, the strand from which the genes are transcribed did not appear to be a determining factor, in contrast to what has been detected for methylation (Patiño-Parrado et al., 2017). As the non-coding genes are poorly annotated in the pig genome, there is a higher risk to annotate a protein-coding gene (adjacent to a not annotated non-coding gene) as being imprinted by error. While we were able to detect the expression of several long non-coding RNAs (COPG2IT1 and AIRN) by visual inspection, *COPG2IT1* and *AIRN* would have been tagged to *COPG2* and *IGF2R*, respectively instead of non-coding genes. These examples showed the importance of genome annotation for studying genomic imprinting in a genome-wide manner. In addition, discrepancies in parental expression along a locus as we observed for the *COPG2* gene need a deep inspection to bring proper conclusion about the imprinting status of a gene. Altogether, the *COPG2* locus demonstrated the complexity and potential for misleading results that can arise from a genome-wide approach.

The first comparative studies regarding the presence of imprinted genes across species and tissues focused on a few candidate genes per orthology, particularly in livestock species (Hubert et al., 2024b), and suggested firstly that these atypical regulatory mechanisms were conserved. High-throughput - omics technologies then provided access to a global and more exhaustive overview, leading to more contrasting results, despite the biases inherent in these methods (Edwards et al., 2023). It was progressively shown that imprinted expression (i.e. the number of imprinted genes as well as the strength of the parental bias for a given gene) was strongest in embryonic tissues including hypothalamus and placenta than in adulthood tissues (Andergassen et al., 2017; Perez et al., 2015; Pinter et al., 2015). More specifically, the hypothalamus has even been characterized with a higher frequency of imprinted gene expression than in other brain regions (Babak et al., 2015; Gregg et al., 2010). Here, results from the comparison between tissues were consistent with these conclusions as we identified 174, 78 and 239 unique genes in the hypothalamus, muscle and placenta, respectively from the least stringent filters. In addition, for the 114 genes showing strongest POE magnitudes, 85% were genes detected in the hypothalamus and placenta supporting the notion that genomic imprinting may constitute a conserved mechanism to instruct both neural and placental functions.

Some discrepancies that we observed across tissues concerning patterns of imprinting has already been shown such as the switch of parental allele expression of *IGF2* in the hypothalamus, which was strongly maternal while it was exclusively expressed from the paternal allele in the other tissues investigated (Gregg et al., 2010; Perez et al., 2015). However, that was not the case for *IGF2R* that has been shown in mice to be imprinted strictly in non-neuronal cells (Perez et al., 2015) while we highlighted, as its ligand *IGF2*, a change in the strength of the parental bias. Furthermore, the placenta is the tissue with the most imprinting discrepancies between the mouse and humans (Monk, 2015) even though the placentas of these two species are structurally equivalent (discoidal or hemo-chorial placenta). From a functional point of view, the dynamics of gene expression during gestation demonstrated that the mouse placenta looked similar to the human placenta only during the first half of pregnancy (Soncin et al., 2018) demonstrating the importance of developmental stage of placenta for comparison. Here, placentas from piglets were sampling around birth likely suggesting specific gene expression profiles. In addition, pig presents a so-called diffuse or epithelo-chorial placenta (Stenhouse et al., 2022) unlike humans and mice. We hypothesized that developmental stage and placentation might explained inconsistencies we observed since we highlighted that the imprinting patterns that were not conserved between species particularly concerned genes expressed in the placenta such as *CBR1*, *HMG13* and *PLAGL1*. In fact, these genes showed exclusive paternal expression in human and murine placentas (Monk, 2015; Monk et al., 2006), unlike in pigs where the opposite was observed.

Our approach revealed a number of genes with a strong or even exclusive parental expression bias (n=114), consistent with studies carried out in model species such as humans and mice in which approximately 200 imprinted have been characterized (Tucci et al., 2019). While genes already functionally known to be imprinted in pigs were detected, such as *NAP1L5* (Jiang et al., 2011), *MEST* (Zhang et al., 2012), *PEG3* (Jiang et al., 2011), novel imprinted genes not yet known to be subjected to genomic imprinting in model organisms were identified such as *POU6F2*, *FAM20B*, *PDE4DIP, HSD17B11 (17-Beta-Hydroxysteroid Dehydrogenase Xi)* and *PITRM1*. Among these genes, it is difficult to determine whether they might be specific to the porcine species or shared to a wider phylogenic clade since some novel imprinted genes still remain to discover (Hu et al., 2023) due to many barriers for their detection including variant informativeness, statistical power and developmental stages. Interestingly, ASE was shown for *PDE4DIP* in both *Bos Indicus* (de Souza et al., 2020) and patients with acute myeloid leukemia (Mulet-Lazaro et al., 2021) likely suggesting mono-allelic expression of this gene in others species but without searching for POE. In addition, these genes appeared to be particularly important for behavioral phenotypes, neural development, growth and pre- or post-natal mortality, as demonstrated by the effects in knockouts mice (Mouse Genome Informatics database, https://www.informatics.jax.org/). For example, deletions of *FAM20B* in mice led to embryonic lethality or perinatal lethality with homozygous embryos showing severely stunted growth, with multisystem organ hypoplasia and delayed development (Liu et al., 2018; Vogel et al., 2012). Furthermore, germline mutations in *POU6F2* were significantly associated with Wilms’ tumor (Perotti et al., 2004), a child nephroblastoma that is highly prevalent in human imprinting disorders such as Beckwith-Wiedemann syndrome (Anvar et al., 2019). Although completely new, it can’t be ruled out that human and murine orthologs of some of these novel imprinted genes have not yet been detected. This could provide a first step towards the identification of other imprinting genes in mammals. This dichotomy in imprinting distribution, gathering on one hand high expression biases represented mostly by known imprinting genes, and on the other hand, moderate to weak parental expression biases represented by a majority of candidate genes reported in the literature, supports the relevance of our approach (Edwards et al., 2023; Perez et al., 2016).

Moreover, both *KBTBD6* (Ahn et al., 2023) and *ZNF791* (Ahn et al., 2025) genes identified from gynogenic embryos as imprinted exclusively in domestic animals including artiodactyls and carnivores are among the genes showing total imbalance in our study. Another gene, with unknown function called *ZNF300-like* (*LOC100520903*) that is currently unannotated in the porcine genome and that does not have an orthologous in the human and murine genomes, showed an exclusive paternal expression (ratio 3 :97) in the placenta and hypothalamus. These results increase the spectrum of imprinted genes that may have acquired their POE during speciation and genome evolution, likely *via* the integration of retroviral sequences (Andergassen et al., 2021). In addition, deeper inspection of specific loci suggesting discrepancies such as the *COPG2* locus highlighted the complexity of data analyses from genome-wide studies since we identified the antisens non-coding *COPG2IT1* gene and its paternal expression in the meantime we showed isoform-specific imprinted for the *COPG2* gene. This specific imprinted pattern observed for the shortest isoforms of COPG2 in hypothalamus might come from a tissue-specific alternative promoter (Stelzer et al., 2015) or might be typical of specific neuronal cells (Gregg et al., 2010; Ho-Shing and Dulac, 2019). However, this novel result never described so far in pigs bring new insights in genomic imprinting mechanisms in pigs although imprinted-isoform-specificity has already been highlighted in different species (Newman et al., 2024; Stelzer et al., 2015).

## Conclusion

The regulatory mechanisms leading to parental expression bias need to be better understood, as has been the case in recent years with the identification of non-canonical mechanisms. In addition, evaluation of the magnitude of genomic imprinting across species, tissues and developmental stages might unravel its origin and implementation of this phenomenon, which is key to essential biological functions such as fetal and post-natal development, growth and neuronal development. Finally, characterization of imprinted genes for livestock species particularly, might significantly improve the evaluation of parental genomes contribution into agronomic complex traits.

## Methods

### Ethical statement

All the procedures and guidelines for animal care were approved by the local ethical committee in animal experimentation (Poitou-Charentes) and the French Ministry of Higher Education and Scientific Research (authorizations n°2018021912005794 and n°11789-2017101117033530).

The use of animals and the procedures performed in this study to obtain placental samples (dataset 2) were approved by the European Union legislation (directive 86/609/EEC) and French legislation in the Midi-Pyrénées Region of France (Decree 2001–464 29/05/01; accreditation for animal housing C-35–275- 32). The technical and scientific staff obtained individual accreditation (MP/01/01/01/11) from the Ethics Committee (région Midi-Pyrénées, France) for experiments involving live. Under these conditions, this study follows the ARRIVE guidelines (Animal Research: Reporting of In Vivo Experiments), and is committed to the 3Rs of laboratory animal research and, thus using the minimum number of animals to achieve statistical significance.

### Animals and experimental designs

In order to detect parental expression bias, we relied on 2 different datasets generated from reciprocal crosses between two genetically distant pig breeds, the European Large White (LW) breed and the Asian Meishan (MS) breed (Fig. 1A). Piglets from the first dataset have been produced to highlight parental expression bias in muscle and hypothalamus at one day by combining transcriptomic data obtained only from piglets and genomic data obtained on both parents and offspring from blood. The second dataset was dedicated to target imprinting genes in the placenta at 110 days of gestation. This developmental stage corresponds to a mature stage in pig since gestation periode ranges between 111 and 120 days from the first day of mating.

Animals from the first reciprocal cross (dataset 1) were generated in the frame of the PIPETTE project (French National Research Agency ANR-18-CE20-0018) described in (Hubert et al., 2024a). Animals were produced in a reciprocal cross design between Large White and Meishan pig breeds. The study included 14 pigs, 13 pigs were bred at the GenESI INRAE experimental farm (https://doi.org/10.15454/1.5572415481185847E12) and 1 pig (Large White boar) came from breeding organizations in accordance with the French and European legislation on animal welfare (Figure 1A). Animals from the second reciprocal cross (dataset 2) were generated in the frame of the PORCINET project (French National Research Agency ANR-09-GENM005). Animals were also produced at the GenESI INRAE experimental farm (https://doi.org/10.15454/1.5572415481185847E12) in a reciprocal cross design between Large White and Meishan pig breeds from 17 sows (10 Large White and 7 Meishan) and 4 boars (2 Large White and 2 Meishan). Details on animal resources and the whole genetic design can also be found in (Lefort et al., 2020) (Fig. 1A).

### Biological sample collection and nucleic acid extraction

For genomic DNA extraction, 14 biological samples were used in the dataset 1. Blood samples were collected on EDTA and were stored frozen nine months at -20°C. Biological samples were collected at adult developmental stage for all the parents (n=5) of the reciprocal cross design while biological samples were collected at 1d after birth for all offspring (n=9) of the reciprocal cross design. High molecular weight genomic DNA was extracted from blood using the Genomic-tip 100/G kit (Qiagen, Reference 10243) as recommended by the manufacturer. Genomic DNA concentrations were measured using the Qubit fluorimetry system with the Broad Range kit (Invitrogen, Reference Q32850) for detection of double-stranded. Fragment size distributions were assessed using the Femto Pulse Genomic DNA 165 kb Kit (Agilent). Purity was measured using a Nanodrop system (Thermo Fisher). All samples were purified with beads to obtain expected ratios i.e. 260/280 : 1.8-2 and 260/230 : 2-2.2.

In addition, genomic DNA was extracted from blood, hypothalamus and muscle (n=4) to perform molecular analyses of the IGF2 locus.

For transcripts RNA extraction from *longissimus dorsi* muscle and hypothalamus from the first reciprocal cross (dataset 1), nine piglets were slaughtered one day after birth. A piece of *longissimus dorsi* and hypothalamus was cut and frozen straightforwardly to avoid degradation of transcripts. RNA extractions for muscle were performed using the Quick-RNA Miniprep Plus Kit (Zymo Research, Reference R1057) following the manufacturer recommendation. RNA extractions for hypothalamus were performed using the NucleoSpin RNA kit (Macherey-Nagel, Reference 740955) following the manufacturer recommendation. RNA concentrations were measured using the Qubit fluorimetry system with the Broad Range kit (Invitrogen, Reference Q10210). Fragment size distributions were assessed using agarose seakem electrophoresis after RNA denaturation. Purity was measured using a Nanodrop 8000 system (Thermo Scientific).

For the second reciprocal cross (dataset 2), the 54 placenta samples and their associated endometrium from crossbred fetuses were collected at 110 days of gestation. RNA extraction was carried out on the 54 frozen powdered placenta samples according to the NucleoSpin RNA, Mini kit for RNA purification (Macherey-Nagel, Reference 740955) Preparatory step, RNA purification and quality controls are described in the protocol submitted to FAANG database (https://api.faang.org/files/protocols/experiments/INRAE_SOP_COLOcATION_RNA_Extraction_20230425.pdf).

### Sequencing technologies

Preparation of all libraries and both RNA and DNA sequencing were performed on the GeT-PlaGe core facility in INRAE Toulouse (https://doi.org/10.17180/NVXJ-5333).

Long reads Oxford Nanopore Technology (ONT) were used to sequence genomic DNA extracted from blood for the dataset 1. ONT libraries were prepared according to the manufacturer’s instructions “1D gDNA selecting for long reads (SQK-LSK109)”. At each step, DNA was quantified using the Qubit dsDNA HS Assay Kit (Life Technologies). DNA purity was tested using the Nanodrop (Thermofisher) and size distribution and degradation assessed using the Fragment Analyzer (Agilent) DNF-464 HS Large Fragment Kit. Purification steps were performed using AMPure XP beads (Beckman Coulter). For one Flowcell, 5μg of DNA were purified then sheared at 25kb using the Megaruptor system (diagenode). A size selection step using the Short Read Eliminator M Kit (Circulomics) was performed. A one step DNA damage repair + END-repair + dA tail of double stranded DNA fragments was performed on 2μg of DNA. Then adapters were ligated to the fragments. The library was loaded onto FLO-PRO002 R9.4.1 flow cell and sequenced on PromethION instrument at 20fmol within 72H. At 24H and 48H a nuclease flush was performed on the same flow cell, the library was re-loaded at 20fmol. The DNAseq of the dataset 1 (blood) are available from the European Nucleotide Archive (ENA) under the accession number PRJEB86771.

Briefly for RNA-seq experiments, libraries were carried out by the GeT-PlaGe core facility in INRAE Toulouse. Sequencing was performed on an Illumina NovaSeq6000 using a paired-end read length of 2x150 pb with the Illumina NovaSeq6000 sequencing kits (S4 flow cell). The RNAseq of the first dataset (muscle and hypothalamus) are available from the European Nucleotide Archive (ENA) under the accession number PRJEB86771. For the second dataset, all the protocols and quality controls concerning the dataset 2 were submitted to FAANG database (https://api.faang.org/files/protocols/experiments/INRAE_SOP_COLOcATION_Library_Preparation_20230509.pdf).

## Data preprocessing

### Variant Calling

All bioinformatic analyses were performed at the genotoul bioinformatics platform Toulouse Occitanie (Bioinfo Genotoul, https://doi.org/10.15454/1.5572369328961167E12) and are presented on Figure 1B.

The ONT (filtered) fastq reads were aligned to the Sscrofa11.1 reference assembly (GCF_000003025.6) using minimap2 with the map_ont flag, resulting in alignments corresponding to a coverage ranging from 30X to 60X of the genome. Variants were called using the PEPPER-Margin-DeepVariant pipeline (https://github.com/kishwarshafin/pepper, (Shafin et al., 2021) using pepper_deepvariant release 0.4 with the ont flag. This model was trained for the guppy 4 basecaller corresponding to the basecaller of the produced ONT reads (MinKNOW-Live-Basecalling version 4.0.11 (flipflop)). Only autosomes have been considered. The resulting gvcf files were merged using GLnexus (https://github.com/dnanexus-rnd/GLnexus, (Lin et al., 2018) resulting in a single vcf file for the 14 animals containing 27,569,029 variants. In order to fill missing genotypes from genomic ONT variant calling, data were imputed using Beagle 4.0 (Browning et al., 2018). This vcf file called Dataset1_DNAseq_variants_final.vcf.gz is deposited in the https://recherche.data.gouv.fr/en public database (https://doi.org/10.57745/RF5PUT).

For RNAseq analyses from *longissimus dorsi* (muscle) and hypothalamus, transcriptome assembly and quantification were performed using the Nextflow nf-core\rnaseq (version nfcore- Nextflow RNA-seq 3.13.2) pipeline. The mapping was launched with STAR software against the Sus scrofa genome reference version 11.1 (Sus_scrofa.Sscrofa11.1.dna.toplevel.fa) and the gene structure annotation version 11.1.109 (Sus_scrofa.Sscrofa11.1.109.gtf. Variant calling from RNAseq has been performed following GATK guidelines and available on https://github.com/gatk-workflows/gatk3-4-rnaseq-germline-snps-indels leading to a single vcf file per tissue (*longissimus dorsi* and hypothalamus) for the 9 animals including 2,204,937 and 3,572,855 variants in *longissimus dorsi* and hypothalamus, respectively. These vcf files called Dataset1_RNAseq_muscle_variants_final.vcf.gz and Dataset1_RNAseq_hypothalamus_variants_final.vcf.gz are deposited in the https://recherche.data.gouv.fr/en public database (https://doi.org/10.57745/RF5PUT).

For the second dataset, we used public data from placenta (fetal tissue) and endometrium (maternal tissue) to reconstruct the parental origin of alleles using imputation and phasing (see below). For RNAseq analyses from placenta and endometrium samples, transcriptome assembly and quantification were performed using the Nextflow nf-core\rnaseq (version nfcore-Nextflow-v20.11.0-edge) pipeline.

The mapping was launched with STAR software against the Sus scrofa genome reference version 11.1 (Sus_scrofa.Sscrofa11.1.dna.toplevel.fa) and the gene structure annotation version 11.1.104 (Sus_scrofa.Sscrofa11.1.104.gtf). Bioinformatic analyses are described in a data paper (Maman-Haddad et al., 2024). The RNAseq fastq files have been deposited in the European Nucleotide Archive (ENA) at EMBL-EBI under accession number PRJEB75252. Variant calling was then performed using a fork of the Nextflow nf-core RNAVAR code from GitHub (dev version, release 104). Two DSL2 modules were integrated into the Nextflow nf-core/RNAVAR pipeline and the haplotype code was modified to add a gVCF generation step. The developments were submitted to Nextflow nf-core RNAVAR manager via a pull request on GitHub and are available on GitHub:https://github.com/SarahMaman/rnavar. VCF indexes for dbSNP and indels were built using tabix-0.2.5 and bcftools-1.14. The STAR indexes (v2.7.5a) and the fasta indexes were built with samtools (v1.19) and GATK CreateSequenceDictionary (v4.4.0.0), based on the references Sus_scrofa.Sscrofa11.1.104.gtf, sus_scrofa.vcf (release-104) and Sus_scrofa.Sscrofa11.1.dna. toplevel.fa. The Nextflow nf-core/RNAVAR pipeline development was performed using default filtering criteria and generated 54 gVCF files from 54 placental samples and 28 gVCF files from 28 endometrium samples. Last, one single vcf file per tissue (placental and endometrium) was generated using GATK GenomicsDBImport (v4.2.6.1). In total, 792,902 and 798,812 variants have been called from placenta and endometrium RNAseq, respectively. The vcf file specific of the placenta called Dataset2_RNAseq_placenta_variants_final.vcf.gz is deposited in the https://recherche.data.gouv.fr/en public database (https://doi.org/10.57745/RF5PUT).

### Imputation and complete variant datasets

Since none whole genome sequencing was performed from the second reciprocal cross (dataset 2), we reconstructed genome representation of both parents and offspring based on endometrium and placenta RNAseq data that represent mother and offspring, respectively as well as a personal genomes database. We used the whole dataset presented in (Lefort et al., 2020; Maman-Haddad et al., 2024) including 256 animals (224 RNAseq placenta from fœtus distributed in 194 trios and 30 duos (4 sires and 28 RNAseq endometrium from dams). Within the 224 fetuses, 112 were sampled at 90 days of gestation while 112 others were sampling at 110 days of gestation. Firstly, based on the scientific question on genomic imprinting that represents a mono-allelic expression dependent of the parent-of-origin, only heterozygous biallelic SNPs were kept from both vcf files of the 54 placental fœtus at 110 days and 28 maternal endometrium using bcftools 1.17 (https://github.com/samtools/bcftools, (Danecek et al., 2021). Secondly, we checked that sibling fetuses had the same maternal genome from endometrium RNAseq variant calling after filtration DP>20 using the Identity By State (IBS) parameter of plink 1.9 (https://github.com/chrchang/plink-ng, (Chang et al., 2015). Since the experimental design was between Large White and Meishan breeds, we used a personal Large White (n=75) and Meishan (n=9) genomes database from 84 pigs to impute with Shapeit5 (https://github.com/odelaneau/shapeit5, (Hofmeister et al., 2023) from RNAseq variants calling to whole genome variant calling. Only autosomes and X chromosome have been considered. Finally, we then checked and filtered out mendelian incompatibilities between mother and fetus using plink 1.9 (https://github.com/chrchang/plink-ng, (Chang et al., 2015). In total, we reconstructed genomic data including 31,311,840 SNPs after pruning, imputation and phasing. This vcf file called Dataset2_DNAseq_variants_final.vcf.gz is deposited in the https://recherche.data.gouv.fr/en public database (https://doi.org/10.57745/RF5PUT).

### Annotation of variants

Annotation of significant variants was performed using both Ensembl annotation release 113 and NCBI annotation release 106 databases since differences are often observed between both especially for the 3’ and 5’ UTR annotation and novel genes identification.

### Detection of parental variants

The SnpSift tool (https://pcingola.github.io/SnpEff/, (Cingolani et al., 2012) was used to generate different allelic combinations on reciprocal crosses trios and duos. Each combination represents a possibility of tracing the parental origin of the alleles transmitted to heterozygous descendants. Thus, for each individual from a trio, 3 combinations enable us to determine the paternal origin of allele 0, and 3 other combinations enable us to determine the maternal origin of allele 0 (Fig. 1B). For duos, only one possible combination per parental origin was used (2 combinations in total) (Fig. 1B). In this way, the combinations were first grouped by parental origin and by individual using the bcftools 1.17 (https://github.com/samtools/bcftools, (Danecek et al., 2021). Each descendant therefore has 2 global parental files. In order to obtain a list of positions that could be used to detect specific parental expression in hypothalamus, muscle and placenta tissues, each genomic parental file obtained either from ONT (hypothalamus and muscle) or reconstructed (placenta) was then cross-referenced with its associated variant file generated from RNAseq experiments. A set of files reporting the common positions between the ONT and RNA-seq data was thus obtained for each parental origin and for each individual. These output files were then merged via bcftools 1.17 (https://github.com/samtools/bcftools, (Danecek et al., 2021) to generate a global file containing, for each descendant of each cross, all the positions exploitable for the detection of POE and ASE. The parental alleles of the individuals’ genotypes were recoded into P (Paternal Origin) and M (Maternal Origin) alleles using a customized pearl script. Consequently, for each variant, two nomenclatures existed, one with alleles coded 0 and 1 to detect ASE and one coded P and M to detect POE.

## Statistical analyses

Classical one-sided Fisher exact tests have been used to detect parental biases expression in each tissue. A first test (ASE test) evaluated for each informative genomic variant (i.e., heterozygous variant in offspring with known parental origin), the presence of allelic bias by comparing the distribution of reads per allele (0 and 1) in RNAseq data to a 50:50 distribution that is typical of biallelic expression. A second test (POE test) evaluated for each informative variant from genomic (i.e., heterozygous variant in offspring with known parental origin), the presence of parental bias by comparing the distribution of reads per parental allele (P and M) in RNAseq data to a 50:50 distribution that is typical of biallelic expression. The two additional tests (ASE *vs.* POE and POE *vs.* ASE tests) evaluated for each informative variant from genomic (i.e. heterozygous variant in offspring with known parental origin), the direction of the bias expression by comparing both minimums and maximums of the distribution of reads given their classification by allele (0 and 1) or parental allele (P and M). A threshold corresponding to a Bonferroni correction (Pval adjusted = number of tests / 0.01) was applied for each statistical test performed (Fig. 1B).

To determine unique parent-of-origin expressed genes, we used only significant annotated variants. For genes that were tagged by several variants, the mean of all maternal expressed alleles was evaluated.

## Molecular analyses of the *IGF2* locus

### Allele-specific PCR for parental informativity of variant

PCR Allele Competitive Extension (PACE) (von Maydell, 2023) analysis was performed with 10 ng of purified blood genomic DNA using the PACE®-IR 2x Genotyping Master mix (3CR Bio-science) in addition of 12 microM of a mix of extended allele specific forward primers and 30 microM of common reverse primer in a final volume of 5 μL. The touch-down PCR amplification condition was 15 min at 94°C for the hot-start (eurobio SCIENTIFIC, reference PB10.11-05) activation, 10 cycles of 20 s at 94°C, 58–66°C for 30 s (dropping 0.8 ◦C per cycle), then 40 cycles of 20 s at 94°C and 30 s at 58°C performed on an ABI9700 thermocycler followed by a final point read of the fluorescence on an ABI QuantStudio 6 real-time PCR system and using the QuantStudio software 1.3 (Applied Biosystems). The trio of primers used for genotyping the variant rs3473951016 are GAAGGTGACCAAGTTCATGCTCCAGGAGCAGAGTGCGCAC, GAAGGTCGGAGTCAACGGATTATCCAGGAGCAGAGTGCGCAA and TCCCTGCCCAGCCCCCACT.

### Bisulfite conversion of genomic DNA extracted from hypothalamus and muscle

500 ng of hypothalamus and muscle genomic DNA sample collected from 10 piglets were bisulfite converted using an EZ DNA methylation-direct kit (Zymo Research, cat D5020).

### PCR conditions before Sanger sequencing

We used Sanger sequencing to (i) validate the accuracy from PACE genotyped rs3473951016 variant from blood genomic DNA samples and (ii) for epigenotyping rs3473951016 variant on sodium bisulfite-treated genomic DNA obtained from hypothalamus and muscle samples.

PCR were performed using the PCR eurobio 5X kit (eurobio SCIENTIFIC, reference PB10.11-05) in the presence of primers at 10 μM, 0.05 U of Taq polymerase and the corresponding amount of genomic DNA. The genotyping step required an amount of 10 ng of blood genomic DNA. The epigenotyping step required 25 ng of bisulfite converted hypothalamus and muscle genomic DNA targeting the IGF2 locus. The pair of primers used to genotype the variant rs3473951016 was CCGGGCTTTTTCTAACAGG and CCGTCGACTAGCTGGTGAAT (genotyping step). The pair of primers used to detect the methylated allele and perform Methylation-sensitive PCR (MS-PCR) was AAAGAGTTTCGTTTTTTTAGGGCGG and AAACCCTAACCCCACACCCCTTACG (epigenotyping step). The PCR cycling conditions were as follows: 5 min at 94°C for the PCRBIO Taq DNA Polymerase Classic activation, then 30 cycles (genotyping) and 35 cycles (epigenotyping) of 30 s at 94°C, primer annealing of 30 s at 60°C, and 30 s of primer extension at 72°C, performed on a verity PCR machine (Applied Biosystems).

The PCR products were then purified using the “Thermosensitive Alkaline Phosphatase” (F0654, Thermo-Scientific) a 0,06 U/microL final), and the “Exonuclease I” (MO293L, NEB) 1,25 U/μM. The reaction was performed at 37°C for 45 min followed by an inactivation step at 80°C during 30min. The sequencing reaction has been realized by Genewiz company (https://www.genewiz.com/en-GB/). The results were analysed with the CLC software.

## Supporting information

Table with sheets for parent-of-origin expressed genes in different tissues

## Declarations

### Ethics approval and consent to participate

Use of animals and the procedures performed in this study were approved by the European Union legislation (directive 86/609/EEC) and French legislation in the Midi-Pyrénées Region of France (Decree 2001–464 29/05/01; accreditation for animal housing C-35–275-32). The technical and scientific staff obtained individual accreditation (MP/01/01/01/11) from the Ethics Committee (région Midi-Pyrénées, France) for experiments involving live. Under these conditions, this study follows the ARRIVE guidelines (Animal Research: Reporting of In Vivo Experiments), and is committed to the 3Rs of laboratory animal research and, thus usingthe minimum number of animals to achieve statistical significance.

## Consent for publication

Not applicable

## Availability of data and materials

The DNAseq and RNASeq fastq files of the dataset 1 (blood, muscle and hypothalamus) are available from the European Nucleotide Archive (ENA) under accession number PRJEB86771. For the dataset 2 (placenta), the RNAseq.fastq files have been deposited in the European Nucleotide Archive (ENA) at EMBL-EBI under accession number PRJEB75252, a datapaper is also available (Maman-Haddad et al., 2024). Preprocessing variant calling files (.vcf) are deposited in the https://recherche.data.gouv.fr/en public database (https://doi.org/10.57745/RF5PUT).

## Authors’ contributions

Mathilde Perret performed bioinformatic and statistical analyses, wrote the manuscript. Nathalie Iannuccelli performed DNA and RNA extractions for hypothalamus and muscle. Sophie Leroux performed sampling and RNA extractions for hypothalamus and muscle. Katia Fève supervised Mathilde Perret for molecular experiments. Patrice Dehais performed pearl script for bioinformatic analyses. Eva Jacomet performed the *IGF2* locus experiments. Jean-Noël Hubert supervised Eva Jacomet. Carole Iampetro performed ONT experiments. Céline Vandecasteele performed sequencing quality criteria. Sarah Maman-Haddad performed preprocessing analyses of RNA sequencing and SNP detection from endometrium and placenta. Thomas Faraut performed variant calling from ONT sequencing. Laurence Liaubet led the project PORCINET (n°ANR-09-GENM005). Agnès Bonnet led the project COLOCATION (n°ANR-20-CE20-0020). Cécile Donnadieu led the SeqOccIn project from the Operational program ERDF-FSE MIDI-PYRENEES ET GARONNE 2014-2020. Juliette Riquet is Mathilde Perret supervisor and analysed the data. Julie Demars led the project PIPETTE (ANR-18-CE20-0018), supervised Mathilde Perret, analysed the data, wrote the manuscript and draw figures. All authors read, edited and approved the manuscript

## Acknowledgements

We greatly thank all the people from the INRAE experimental unit (GenESI, https://doi.org/10.15454/1.5572415481185847E12) for taking care of animals. We greatly thank all the people from the bioinformatics core facility at INRAE Toulouse (Bioinfo Genotoul, https://doi.org/10.15454/1.5572369328961167E12) for their continuous and efficient support. We greatly thank all the people from the genomics core facility at INRAE Toulouse (GeT-PlaGe Genotoul, https://doi.org/10.17180/NVXJ-5333) for their skills in various sequencing technologies.

## References

Ahn, J., Hwang, I.-S., Park, M.-R., Cho, I.-C., Hwang, S., Lee, K., 2020. The Landscape of Genomic Imprinting at the Porcine SGCE/PEG10 Locus from Methylome and Transcriptome of Parthenogenetic Embryos. G3 GenesGenomesGenetics 10, 4037–4047. 10.1534/g3.120.401425

Ahn, J., Hwang, I.-S., Park, M.-R., Hwang, S., Lee, K., 2023. Imprinting at the KBTBD6 locus involves species-specific maternal methylation and monoallelic expression in livestock animals. J. Anim. Sci. Biotechnol. 14, 131. 10.1186/s40104-023-00931-3

Ahn, J., Hwang, I.-S., Park, M.-R., Rosa-Velazquez, M., Cho, I.-C., Relling, A.E., Hwang, S., Lee, K., 2025. Evolutionary lineage-specific genomic imprinting at the ZNF791 locus. PLOS Genet. 21, e1011532. 10.1371/journal.pgen.1011532

Andergassen, D., Dotter, C.P., Wenzel, D., Sigl, V., Bammer, P.C., Muckenhuber, M., Mayer, D., Kulinski, T.M., Theussl, H.-C., Penninger, J.M., Bock, C., Barlow, D.P., Pauler, F.M., Hudson, Q.J., 2017. Mapping the mouse Allelome reveals tissue-specific regulation of allelic expression. eLife 6, e25125. 10.7554/eLife.25125

Andergassen, D., Smith, Z.D., Kretzmer, H., Rinn, J.L., Meissner, A., 2021. Diverse epigenetic mechanisms maintain parental imprints within the embryonic and extraembryonic lineages. Dev. Cell 56, 2995–3005.e4. 10.1016/j.devcel.2021.10.010

Anvar, Z., Acurzio, B., Roma, J., Cerrato, F., Verde, G., 2019. Origins of DNA methylation defects in Wilms tumors. Cancer Lett. 457, 119–128. 10.1016/j.canlet.2019.05.013

Babak, T., DeVeale, B., Tsang, E.K., Zhou, Y., Li, X., Smith, K.S., Kukurba, K.R., Zhang, R., Li, J.B., Van Der Kooy, D., Montgomery, S.B., Fraser, H.B., 2015. Genetic conflict reflected in tissue-specific maps of genomic imprinting in human and mouse. Nat. Genet. 47, 544–549. 10.1038/ng.3274

Bischoff, S.R., Tsai, S., Hardison, N., Motsinger-Reif, A.A., Freking, B.A., Nonneman, D., Rohrer, G., Piedrahita, J.A., 2009. Characterization of conserved and nonconserved imprinted genes in swine. Biol. Reprod. 81, 906–920. 10.1095/biolreprod.109.078139

Bonthuis, P.J., Steinwand, S., Stacher Hörndli, C.N., Emery, J., Huang, W.-C., Kravitz, S., Ferris, E., Gregg, C., 2022. Noncanonical genomic imprinting in the monoamine system determines naturalistic foraging and brain-adrenal axis functions. Cell Rep. 38, 110500. 10.1016/j.celrep.2022.110500

Browning, B.L., Zhou, Y., Browning, S.R., 2018. A One-Penny Imputed Genome from Next-Generation Reference Panels. Am. J. Hum. Genet. 103, 338–348. 10.1016/j.ajhg.2018.07.015

Chang, C.C., Chow, C.C., Tellier, L.C., Vattikuti, S., Purcell, S.M., Lee, J.J., 2015. Second-generation PLINK: rising to the challenge of larger and richer datasets. GigaScience 4, s13742-015–0047–8. 10.1186/s13742-015-0047-8

Cingolani, P., Patel, V.M., Coon, M., Nguyen, T., Land, S.J., Ruden, D.M., Lu, X., 2012. Using Drosophila melanogaster as a Model for Genotoxic Chemical Mutational Studies with a New Program, SnpSift. Front. Genet. 3. 10.3389/fgene.2012.00035

Danecek, P., Bonfield, J.K., Liddle, J., Marshall, J., Ohan, V., Pollard, M.O., Whitwham, A., Keane, T., McCarthy, S.A., Davies, R.M., Li, H., 2021. Twelve years of SAMtools and BCFtools. GigaScience 10, giab008. 10.1093/gigascience/giab008

Daskeviciute, D., Chappell-Maor, L., Sainty, B., Arnaud, P., Iglesias-Platas, I., Simon, C., Okae, H., Arima, T., Vassena, R., Lartey, J., Monk, D., 2025. Non-canonical imprinting, manifesting as post-fertilization placenta-specific parent-of-origin dependent methylation, is not conserved in humans. Hum. Mol. Genet. ddaf009. 10.1093/hmg/ddaf009

de Souza, M.M., Zerlotini, A., Rocha, M.I.P., Bruscadin, J.J., Diniz, W.J. da S., Cardoso, T.F., Cesar, A.S.M., Afonso, J., Andrade, B.G.N., Mudadu, M. de A., Mokry, F.B., Tizioto, P.C., de Oliveira, P.S.N., Niciura, S.C.M., Coutinho, L.L., Regitano, L.C. de A., 2020. Allele-specific expression is widespread in Bos indicus muscle and affects meat quality candidate genes. Sci. Rep. 10, 10204. 10.1038/s41598-020-67089-0

Edwards, C.A., Takahashi, N., Corish, J.A., Ferguson-Smith, A.C., 2019. The origins of genomic imprinting in mammals. Reprod. Fertil. Dev. 31, 1203–1218. 10.1071/RD18176

Edwards, C.A., Watkinson, W.M., Telerman, S.B., Hulsmann, L.C., Hamilton, R.S., Ferguson-Smith, A.C., 2023. Reassessment of weak parent-of-origin expression bias shows it rarely exists outside of known imprinted regions. eLife 12, e83364. 10.7554/eLife.83364

Gregg, C., Zhang, J., Weissbourd, B., Luo, S., Schroth, G.P., Haig, D., Dulac, C., 2010. High-resolution analysis of parent-of-origin allelic expression in the mouse brain. Science 329, 643–648. 10.1126/science.1190830

Hofmeister, R.J., Ribeiro, D.M., Rubinacci, S., Delaneau, O., 2023. Accurate rare variant phasing of whole-genome and whole-exome sequencing data in the UK Biobank. Nat. Genet. 55, 1243–1249. 10.1038/s41588-023-01415-w

Ho-Shing, O., Dulac, C., 2019. Influences of genomic imprinting on brain function and behavior. Curr. Opin. Behav. Sci., Genetic Imprinting and behaviour (2019) 25, 66–76. 10.1016/j.cobeha.2018.08.008

Hu, Y., Yuan, S., Du, X., Liu, J., Zhou, W., Wei, F., 2023. Comparative analysis reveals epigenomic evolution related to species traits and genomic imprinting in mammals. The Innovation 4, 100434. 10.1016/j.xinn.2023.100434

Hubert, J.-N., Iannuccelli, N., Cabau, C., Jacomet, E., Billon, Y., Serre, R.-F., Vandecasteele, C., Donnadieu, C., Demars, J., 2024a. Detection of DNA methylation signatures through the lens of genomic imprinting. Sci. Rep. 14, 1694. 10.1038/s41598-024-52114-3

Hubert, J.-N., Perret, M., Riquet, J., Demars, J., 2024b. Livestock species as emerging models for genomic imprinting. Front. Cell Dev. Biol. 12, 1348036. 10.3389/fcell.2024.1348036

Inoue, K., Hirose, M., Inoue, H., Hatanaka, Y., Honda, A., Hasegawa, A., Mochida, K., Ogura, A., 2017. The Rodent-Specific MicroRNA Cluster within the Sfmbt2 Gene Is Imprinted and Essential for Placental Development. Cell Rep. 19, 949–956. 10.1016/j.celrep.2017.04.018

Ishihara, T., Suzuki, S., Newman, T.A., Fenelon, J.C., Griffith, O.W., Shaw, G., Renfree, M.B., 2024. Marsupials have monoallelic MEST expression with a conserved antisense lncRNA but MEST is not imprinted. Heredity 132, 5–17. 10.1038/s41437-023-00656-z

Jiang, C.D., Li, S., Deng, C.Y., 2011. Assessment of genomic imprinting of PPP1R9A, NAP1L5 and PEG3 in pigs. Genetika 47, 537–542.

Lefort, G., Servien, R., Quesnel, H., Billon, Y., Canario, L., Iannuccelli, N., Canlet, C., Paris, A., Vialaneix, N., Liaubet, L., 2020. The maturity in fetal pigs using a multi-fluid metabolomic approach. Sci. Rep. 10, 19912. 10.1038/s41598-020-76709-8

Lin, M.F., Rodeh, O., Penn, J., Bai, X., Reid, J.G., Krasheninina, O., Salerno, W.J., 2018. GLnexus: joint variant calling for large cohort sequencing. 10.1101/343970

Liu, X., Li, N., Zhang, H., Liu, J., Zhou, N., Ran, C., Chen, X., Lu, Y., Wang, X., Qin, C., Xiao, J., Liu, C., 2018. Inactivation of Fam20b in the neural crest-derived mesenchyme of mouse causes multiple craniofacial defects. Eur. J. Oral Sci. 126, 433–436. 10.1111/eos.12563

Magee, D.A., Berkowicz, E.W., Sikora, K.M., Berry, D.P., Park, S.D.E., Kelly, A.K., Sweeney, T., Kenny, D.A., Evans, R.D., Wickham, B.W., Spillane, C., Machugh, D.E., 2010. A catalogue of validated single nucleotide polymorphisms in bovine orthologs of mammalian imprinted genes and associations with beef production traits. Anim. Int. J. Anim. Biosci. 4, 1958–1970. 10.1017/S1751731110001163

Maman-Haddad, S., Gress, L., Suin, A., Vialaneix, N., Bonnet, A., 2024. RNA-seq data of pig placenta and endometrium during late gestation. Data Brief 57, 111178. 10.1016/j.dib.2024.111178

Mattick, J.S., Amaral, P.P., Carninci, P., Carpenter, S., Chang, H.Y., Chen, L.-L., Chen, R., Dean, C., Dinger, M.E., Fitzgerald, K.A., Gingeras, T.R., Guttman, M., Hirose, T., Huarte, M., Johnson, R., Kanduri, C., Kapranov, P., Lawrence, J.B., Lee, J.T., Mendell, J.T., Mercer, T.R., Moore, K.J., Nakagawa, S., Rinn, J.L., Spector, D.L., Ulitsky, I., Wan, Y., Wilusz, J.E., Wu, M., 2023. Long non-coding RNAs: definitions, functions, challenges and recommendations. Nat. Rev. Mol. Cell Biol. 24, 430–447. 10.1038/s41580-022-00566-8

Monk, D., 2015. Genomic imprinting in the human placenta. Am. J. Obstet. Gynecol. 213, S152–162. 10.1016/j.ajog.2015.06.032

Monk, D., Arnaud, P., Apostolidou, S., Hills, F.A., Kelsey, G., Stanier, P., Feil, R., Moore, G.E., 2006. Limited evolutionary conservation of imprinting in the human placenta. Proc. Natl. Acad. Sci. U. S. A. 103, 6623–6628. 10.1073/pnas.0511031103

Monk, D., Mackay, D.J.G., Eggermann, T., Maher, E.R., Riccio, A., 2019. Genomic imprinting disorders: lessons on how genome, epigenome and environment interact. Nat. Rev. Genet. 20, 235–248. 10.1038/s41576-018-0092-0

Mulet-Lazaro, R., van Herk, S., Erpelinck, C., Bindels, E., Sanders, M.A., Vermeulen, C., Renkens, I., Valk, P., Melnick, A.M., de Ridder, J., Rehli, M., Gebhard, C., Delwel, R., Wouters, B.J., 2021. Allele-specific expression of *GATA2* due to epigenetic dysregulation in *CEBPA* double-mutant AML. Blood 138, 160–177. 10.1182/blood.2020009244

Newman, T., Bond, D.M., Ishihara, T., Rizzoli, P., Gouil, Q., Hore, T.A., Shaw, G., Renfree, M.B., 2024. PRKACB is a novel imprinted gene in marsupials. Epigenetics Chromatin 17, 29. 10.1186/s13072-024-00552-8

Oczkowicz, M., Szmatoła, T., Piórkowska, K., Ropka-Molik, K., 2018. Variant calling from RNA-seq data of the brain transcriptome of pigs and its application for allele-specific expression and imprinting analysis. Gene 641, 367–375. 10.1016/j.gene.2017.10.076

Park, C.-H., Uh, K.-J., Mulligan, B.P., Jeung, E.-B., Hyun, S.-H., Shin, T., Ka, H., Lee, C.-K., 2011. Analysis of imprinted gene expression in normal fertilized and uniparental preimplantation porcine embryos. PloS One 6, e22216. 10.1371/journal.pone.0022216

Patiño-Parrado, I., Gómez-Jiménez, Á., López-Sánchez, N., Frade, J.M., 2017. Strand-specific CpG hemimethylation, a novel epigenetic modification functional for genomic imprinting. Nucleic Acids Res. 45, 8822–8834. 10.1093/nar/gkx518

Perez, J.D., Rubinstein, N.D., Dulac, C., 2016. New Perspectives on Genomic Imprinting, an Essential and Multifaceted Mode of Epigenetic Control in the Developing and Adult Brain. Annu. Rev. Neurosci. 39, 347–384. 10.1146/annurev-neuro-061010-113708

Perez, J.D., Rubinstein, N.D., Fernandez, D.E., Santoro, S.W., Needleman, L.A., Ho-Shing, O., Choi, J.J., Zirlinger, M., Chen, S.-K., Liu, J.S., Dulac, C., 2015. Quantitative and functional interrogation of parent-of-origin allelic expression biases in the brain. eLife 4, e07860. 10.7554/eLife.07860

Perotti, D., De Vecchi, G., Testi, M.A., Lualdi, E., Modena, P., Mondini, P., Ravagnani, F., Collini, P., Di Renzo, F., Spreafico, F., Terenziani, M., Sozzi, G., Fossati-Bellani, F., Radice, P., 2004. Germline mutations of the POU6F2 gene in Wilms tumors with loss of heterozygosity on chromosome 7p14. Hum. Mutat. 24, 400–407. 10.1002/humu.20096

Pertea, M., Pertea, G.M., Antonescu, C.M., Chang, T.-C., Mendell, J.T., Salzberg, S.L., 2015. StringTie enables improved reconstruction of a transcriptome from RNA-seq reads. Nat. Biotechnol. 33, 290–295. 10.1038/nbt.3122

Pinter, S.F., Colognori, D., Beliveau, B.J., Sadreyev, R.I., Payer, B., Yildirim, E., Wu, C., Lee, J.T., 2015. Allelic Imbalance Is a Prevalent and Tissue-Specific Feature of the Mouse Transcriptome. Genetics 200, 537–549. 10.1534/genetics.115.176263

Quan, J., Yang, M., Wang, X., Cai, G., Ding, R., Zhuang, Z., Zhou, S., Tan, S., Ruan, D., Wu, Jiajin, Zheng, E., Zhang, Z., Liu, L., Meng, F., Wu, Jie, Xu, C., Qiu, Y., Wang, S., Lin, M., Li, S., Ye, Y., Zhou, F., Lin, D., Li, X., Deng, S., Zhang, Y., Yao, Z., Gao, X., Yang, Y., Liu, Y., Zhan, Y., Liu, Z., Zhang, J., Ma, F., Yang, Jifei, Chen, Q., Yang, Jisheng, Ye, J., Dong, L., Gu, T., Huang, S., Xu, Z., Li, Z., Yang, Jie, Huang, W., Wu, Z., 2024. Multi-omic characterization of allele-specific regulatory variation in hybrid pigs. Nat. Commun. 15, 5587. 10.1038/s41467-024-49923-5

Richard Albert, J., Kobayashi, T., Inoue, A., Monteagudo-Sánchez, A., Kumamoto, S., Takashima, T., Miura, A., Oikawa, M., Miura, F., Takada, S., Hirabayashi, M., Korthauer, K., Kurimoto, K., Greenberg, M.V.C., Lorincz, M., Kobayashi, H., 2023. Conservation and divergence of canonical and non-canonical imprinting in murids. Genome Biol. 24, 48. 10.1186/s13059-023-02869-1

Robinson, J.T., Thorvaldsdottir, H., Turner, D., Mesirov, J.P., 2023. igv.js: an embeddable JavaScript implementation of the Integrative Genomics Viewer (IGV). Bioinformatics 39, btac830. 10.1093/bioinformatics/btac830

Shafin, K., Pesout, T., Chang, P.-C., Nattestad, M., Kolesnikov, A., Goel, S., Baid, G., Kolmogorov, M., Eizenga, J.M., Miga, K.H., Carnevali, P., Jain, M., Carroll, A., Paten, B., 2021. Haplotype-aware variant calling with PEPPER-Margin-DeepVariant enables high accuracy in nanopore long-reads. Nat. Methods 18, 1322–1332. 10.1038/s41592-021-01299-w

Soncin, F., Khater, M., To, C., Pizzo, D., Farah, O., Wakeland, A., Arul Nambi Rajan, K., Nelson, K.K., Chang, C.-W., Moretto-Zita, M., Natale, D.R., Laurent, L.C., Parast, M.M., 2018. Comparative analysis of mouse and human placentae across gestation reveals species-specific regulators of placental development. Development 145, dev156273. 10.1242/dev.156273

Stelzer, Y., Bar, S., Bartok, O., Afik, S., Ronen, D., Kadener, S., Benvenisty, N., 2015. Differentiation of human parthenogenetic pluripotent stem cells reveals multiple tissue- and isoform-specific imprinted transcripts. Cell Rep. 11, 308–320. 10.1016/j.celrep.2015.03.023

Stenhouse, C., Seo, H., Wu, G., Johnson, G.A., Bazer, F.W., 2022. Insights into the Regulation of Implantation and Placentation in Humans, Rodents, Sheep, and Pigs. Adv. Exp. Med. Biol. 1354, 25–48. 10.1007/978-3-030-85686-1_2

Stringer, J.M., Pask, A.J., Shaw, G., Renfree, M.B., 2014. Post-natal imprinting: evidence from marsupials. Heredity 113, 145–155. 10.1038/hdy.2014.10

Tucci, V., Isles, A.R., Kelsey, G., Ferguson-Smith, A.C., Tucci, V., Bartolomei, M.S., Benvenisty, N., Bourc’his, D., Charalambous, M., Dulac, C., Feil, R., Glaser, J., Huelsmann, L., John, R.M., McNamara, G.I., Moorwood, K., Muscatelli, F., Sasaki, H., Strassmann, B.I., Vincenz, C., Wilkins, J., Isles, A.R., Kelsey, G., Ferguson-Smith, A.C., 2019. Genomic Imprinting and Physiological Processes in Mammals. Cell 176, 952–965. 10.1016/j.cell.2019.01.043

Vogel, P., Hansen, G.M., Read, R.W., Vance, R.B., Thiel, M., Liu, J., Wronski, T.J., Smith, D.D., Jeter-Jones, S., Brommage, R., 2012. Amelogenesis imperfecta and other biomineralization defects in Fam20a and Fam20c null mice. Vet. Pathol. 49, 998–1017. 10.1177/0300985812453177

von Maydell, D., 2023. PCR Allele Competitive Extension (PACE). Methods Mol. Biol. Clifton NJ 2638, 263–271. 10.1007/978-1-0716-3024-2_18

Wolf, J.B., Oakey, R.J., Feil, R., 2014. Imprinted gene expression in hybrids: perturbed mechanisms and evolutionary implications. Heredity 113, 167–175. 10.1038/hdy.2014.11

Wu, Y.-Q., Zhao, H., Li, Y.-J., Khederzadeh, S., Wei, H.-J., Zhou, Z.-Y., Zhang, Y.-P., 2020. Genome-wide identification of imprinted genes in pigs and their different imprinting status compared with other mammals. Zool. Res. 41, 721–725. 10.24272/j.issn.2095-8137.2020.072

Zhang, F.W., Han, Z.B., Deng, C.Y., He, H.J., Wu, Q., 2012. Conservation of genomic imprinting at the NDN, MAGEL2 and MEST loci in pigs. Genes Genet. Syst. 87, 53–58. 10.1266/ggs.87.53

