## Additional figures for "An extensive and unbiased genome-wide scan for parent-of-origin expressed genes in the pig clarifies the conservation landscape of genomic imprinting"

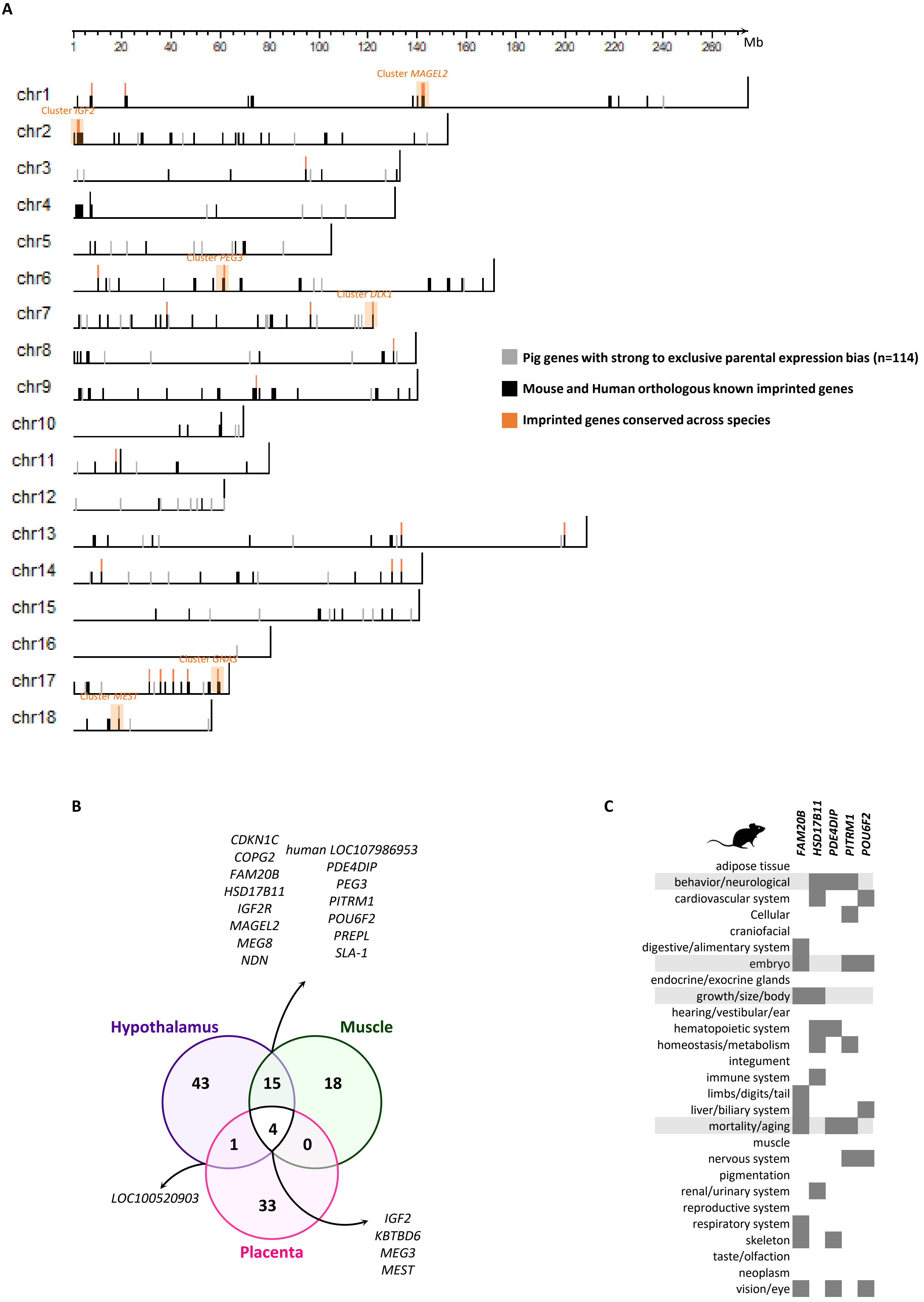


**Additional file 2**

**Additional file 2: Genomic imprinting conservation between species**

**A.** The porcine genome ideogram (autosomes) represents the distribution of genes with high parental expression bias (above 25:75) in the porcine hypothalamus, muscle and placenta, representing a total of 114 unique genes (grey bars). Human and murine known imprinted orthologs are also represented (black bars). Imprinting-conserved genes between pigs and known human and mouse is highlighted in orange. **B.** Venn diagram represents the relations between the 114 unique genes with high parental expression bias in hypothalamus, muscle and placenta. **C.** Novel imprinted genes involved in neurodevelopmental and fetal functions. A set of 5 common imprinting candidates between the hypothalamus and the muscle with strong parental expression bias were identified: *FAM20B, HSD17B11, PDE4DIP, PITRM1, and POU6F2*. We searched for phenotypes associated with these genes (dark grey squares) via the HMDC database: https://www.informatics.jax.org/diseasePortal. Among the identified phenotypes, 4 represent major phenotypes regulated by imprinted genes in mammals: Behaviour/neurological; Embryo; Growth/size/body; Motality/aging (highlighted in light grey).

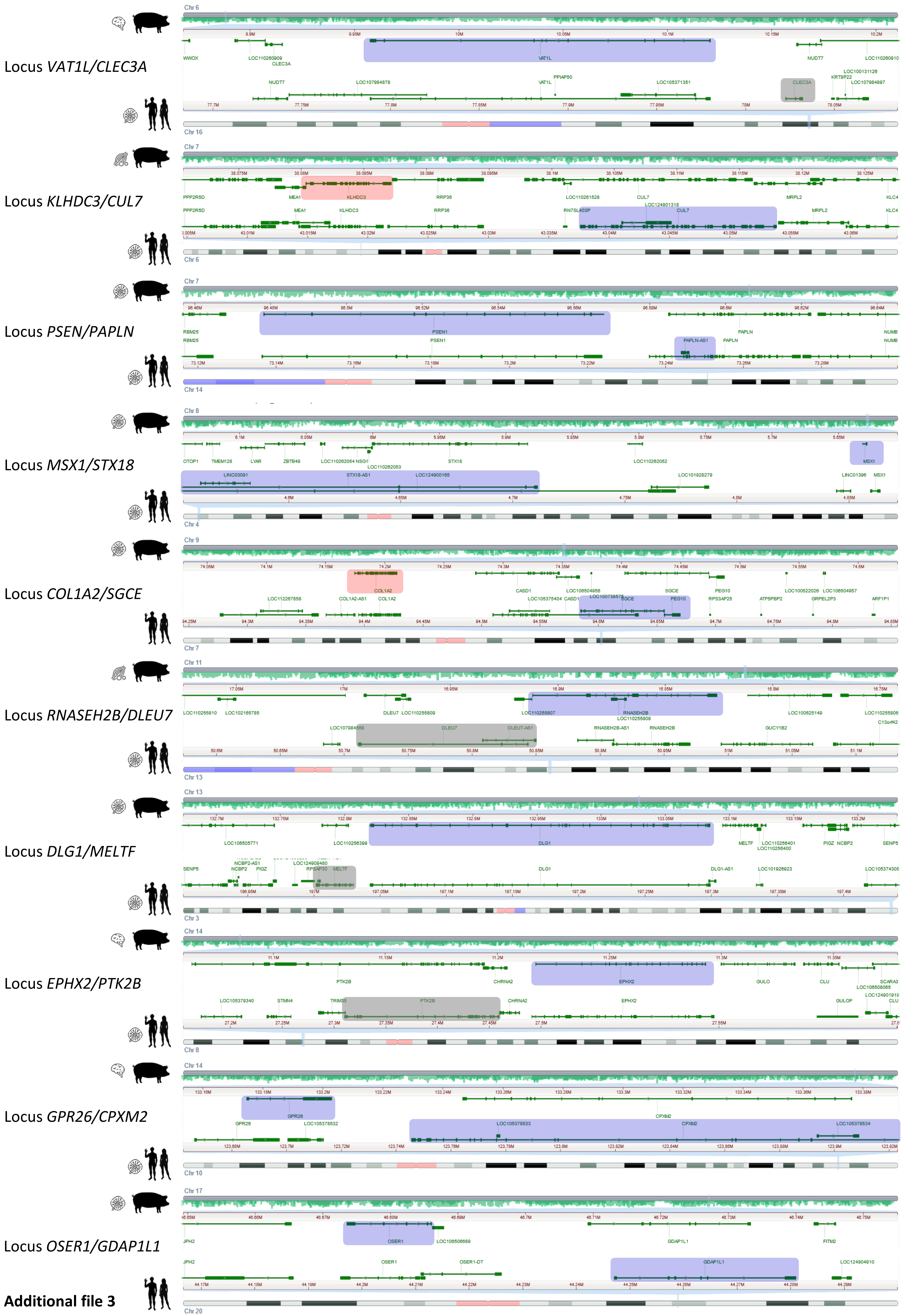


**Additional file 3**

**Additional file 3: Syntenic comparison of imprinted loci identified in pigs and humans**. Overview of regions surrounding 10 genes showing a strong and significant parental expression bias (paternal in blue and maternal in red) in pigs in hypothalamus, muscle or placenta as symbolized by iconography. For the given genes identified including *VATL1, KLHDC3, PSEN, MSX1, COL1A2, RNASEH2B, DLG1, EPHX2, GPR26, OSER1*, a neighboring imprinted gene exists (mostly with a paternal expression (blue) or an undetermined parental expression (grey)) in human tissues.

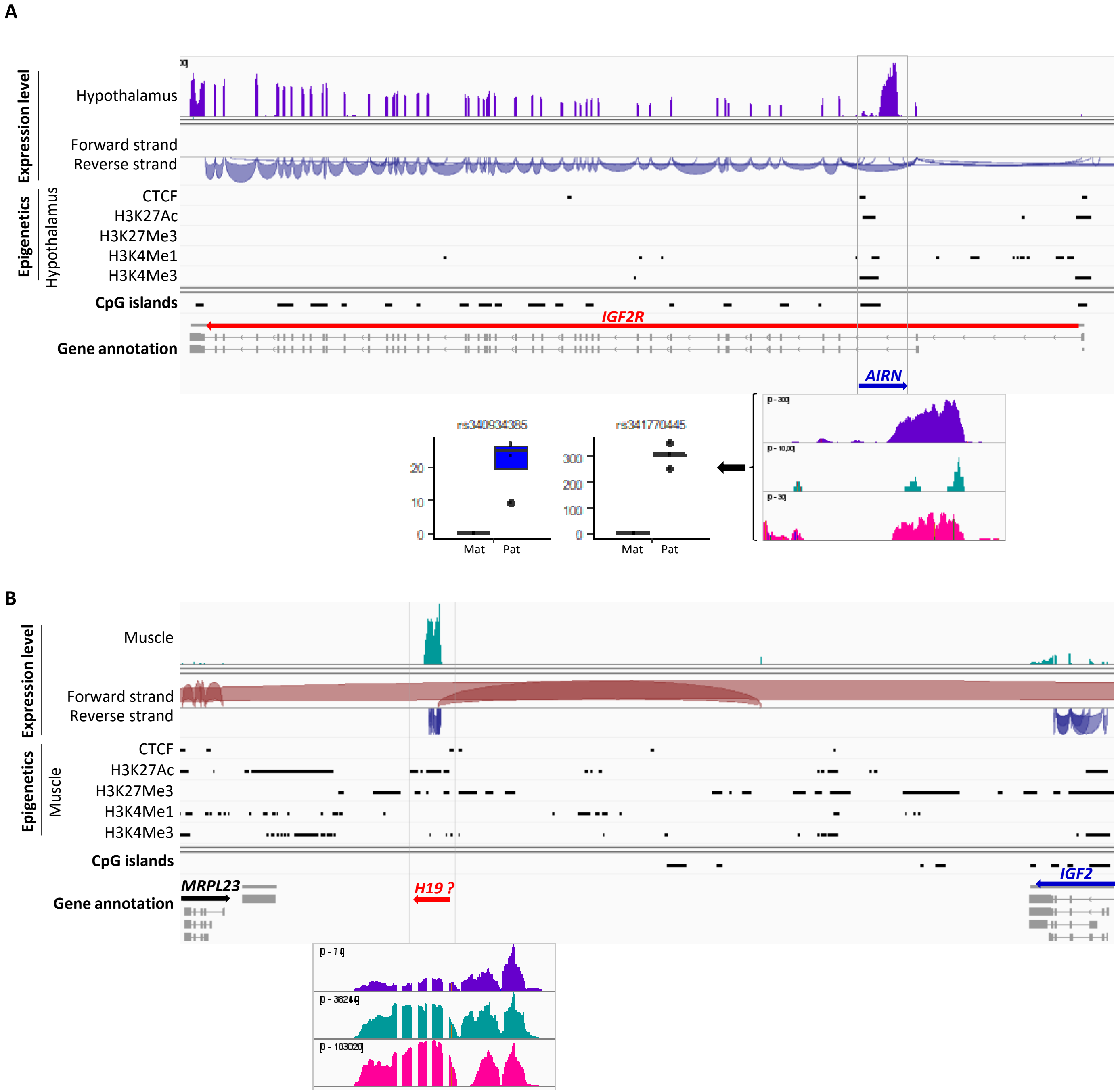


Additional file 4

**<u>Additional file 4:</u> Manual detection of imprinted non-coding genes**

**A.** AIRN identification. RNAseq data show an increased expression in a region of the *IGF2R* gene (grey box). Epigenetic marks, from publicaly available FAANG dataset (https://data.faang.org/home), in this region demonstrate typical marks of genomic imprinting such as H3K4me3. Taken together, these data testify to the presence of an active unannotated lncRNA within in the porcine genome, AIRN. Furthermore, two informative variants rs340934385 and rs341770445 located within this new region showed a trend of paternal expression bias (boxplot). **B**. H19 identification**.** RNAseq data show a strong expression in the unannotated intergenic region of the porcine genome located downstream of the *IGF2* gene (grey box). This region contains typical marks of genomic imprinting such as H3K4me3.. Although highly expressed in the three tissues hypothalamus (purple), muscle (green) and placenta (pink), no informative variants were identified to characterize a parental expression bias.
